# Site-specific structure and stability constrained substitution models improve phylogenetic inference

**DOI:** 10.1101/2023.01.22.525075

**Authors:** Ivan Lorca-Alonso, Miguel Arenas, Ugo Bastolla

## Abstract

In previous studies, we presented site-specific substitution models of protein evolution based on selection on the folding stability of the native state (Stab-CPE), which predict more realistically the evolutionary variability across protein sites. However, those Stab-CPE present qualitative differences from observed data, probably because they ignore changes in the native structure, despite empirical studies suggesting that conservation of the native structure is a stronger selective force than selection on folding stability.

Here we present novel structurally constrained substitution models (Str-CPE) based on Julián Echave’s model of the structural change due to a mutation as the linear response of the protein to a perturbation and on the explicit model of the perturbation generated by a specific amino-acid mutation. Compared to our previous Stab-CPE models, the novel Str-CPE models are more stringent (they predict lower sequence entropy and substitution rate), provide higher likelihood to multiple sequence alignments (MSA) that include one or more known structures, and better predict the observed conservation across sites. The models that combine Str-CPE and Stab-CPE models are even more stringent and fit the empirical MSAs better. We refer collectively to our models as structure and stability constrained substitution models (SSCPE). Importantly in comparison to the traditional empirical substitution models, the SSCPE models infer phylogenetic trees of distantly related proteins more similar to reference trees based on structural information. We implemented the SSCPE models in the program SSCPE.pl, freely available at https://github.com/ugobas/SSCPE, which infers phylogenetic trees under the SSCPE models with the program RAxML-NG from a concatenated alignment and a list of protein structures that overlap with it.

---

Substitution models of protein evolution are a key element of probabilistic phylogenetic inference. However, they necessarily trade-off realism for computational treatability. In particular, traditional empirical substitution models assume that protein sites evolve independently of each other and under the same substitution process (i.e. frequencies and exchangeabilities are the same), although these assumptions are clearly violated by the physical interactions among amino acids that underlie protein structural integrity and functional dynamics (Liberles et al. 2012, Wilke 2012, Sikosek and Chan 2014). In particular, buried sites are much proner to be ocupied by hydrophobic amino acids, and the substitution rate varies substantially across protein sites, with sites that are exposed in the native state varying much more rapidly than buried sites (Franzosa and Xia 2009, Echave et al. 2016). The failure to take into account this variability across sites may affect the accuracy of branch lengths estimated through substitution models (Yang, 1996; Buckley et al. 2001), which are essential ingredients of phylogenetic inference both in the context of distance-based methods such as Neighbor Joining (Saitou and Nei 1987) and of probabilistic methods such as maximum likelihood (ML) (Felsenstein 1981) and Bayesian inference (Yang et al. 1997). For modelling substitution rate variation across sites, Yang (1993, 1996) proposed to adopt a gamma distribution (+G), fitting its shape parameter from the data. This approach greatly improves the fit of the data, but still it assumes the same substitution process at each site. Another approach to model rate variation was proposed by Halpern and Bruno (1998). Given a site where natural selection produces specific patterns of amino acid frequencies, they proposed how to derive an exchangeability matrix that represents the combined effect of mutation and the Fisher’s fixation probability (Fisher 1930). This approach was later extended by Yang and Nielsen (2008) to selection for codon usage. However, this approach obtains the site-specific frequencies from the data, with the risk of substantial overfitting, instead of modelling how the substitution process is influenced by the protein structure.

More recently, several groups proposed biophysical models that predict how the site-specific evolutionary properties, such as the sequence variability and substitution rate, depend on the structural properties of the site in the native state, i.e. the biologically active state in which the protein has a well-defined average folded conformation (except for intrinsically disordered proteins, see Uversky 2011). These models belong to two main classes.

(1) Stability constrained models of protein evolution (Stab-CPE) represent fitness as the probability to find the protein in the native state (Goldstein 2011, Serohijos and Shakhnovich 2014, Bastolla et al. 2017), which can be estimated from the predicted free energy difference Δ*G* between the native and non-native states of the protein. For computational simplicity, these models assume that only the folding free energy changes upon mutation, while the average structure of the native state remains fixed.

(2) Structurally constrained models of protein evolution (Str-CPE) consider fitness as a decreasing function of the deviation of the average structure of the native state from a reference structure available in the Protein Data Bank (PDB) (Echave 2008). These models predict this deviation through linear response theory applied to the Elastic Network Model (ENM; Tirion 1996, Atilgan et al. 2001) that approximates the native ensemble of the protein, and they assume that the folding free energy remains constant. Although Str-CPE models were originally referenced as stability constrained models, we recommend that it is important to differentiate their name from strict Stab-CPE because they are based on different assumptions.

Some time ago, we developed Stab-CPE models that compute site-specific substitution processes that can be used in phylogenetic inference based on a simplified model of the stability of the native state against both unfolded and misfolded states (Bastolla et al. 2006, Arenas et al. 2015). These substitution models retain high computational efficiency due to the approximation that different sites evolve independently, and they only fit from the data one global parameter that represents the strength of natural selection and 19 global amino acid frequencies as in the +F empirical substitution models of protein evolution. Our Stab-CPE models assigned better AIC and BIC scores to real multiple sequence alignments than empirical substitution models that are the same at all positions (Arenas et al. 2015, Arenas et al. 2019).

Computer simulations based on Stab-CPE and Str-CPE models showed that both models predict that the sites that form more native contacts, which are buried in the native state, are more constrained by negative selection and tend to evolve more slowly, which is consistent with observations from empirical data (Echave et al. 2015). This occurs because mutations at sites with many contacts strongly affect both protein folding stability and the overall protein structure of the native state. However, the Stab-CPE model adopted in these simulations did not consider the stability of the native state against incorrectly folded (misfolded) conformations. Previous studies showed that stability against misfolding is an important selective force in protein evolution, whose footprint can be identified in natural protein structures (Berezovsky et al. 2007, Noivirt-Brik et al. 2009, Minning et al. 2013). Consistently, Stab-CPE models that neglect misfolding are worse supported by empirical data, since they assign lower likelihood to real protein sequences and they predict protein sequences that are on the average more hydrophobic than natural proteins and are not stable against misfolding (since hydrophobicity stabilizes compact conformations belonging to both the native and misfolded states), while models that consider stability against misfolding score better under all these points of view (Arenas et al. 2015). The Stab-CPE models that consider misfolding predict that the protein sites that are more variable and evolve more rapidly are amphiphilic sites with an intermediate number of contacts, which can host both hydrophobic and polar residues (Jimenez et al. 2018). This prediction contrasts with the observation that the most variable sites are exposed sites, and supports Str-CPE against Stab-CPE.

Str-CPE models are also supported by studies that suggest that protein structure is more conserved than protein sequence when the protein function is conserved (Illergard et al. 2009, Pascual-Garcia et al. 2010). This implies that mutations of the protein sequence that do not modify the structure are more likely tolerated, which suggests that negative selection targets protein structure more strongly than protein sequence. Nevertheless, pairs of proteins with different molecular function show variations of the substitution rate attributable to positive selection. These evolutionary accelerations are stronger for protein structure changes than protein sequence changes (Pascual-Garcia et al. 2019), again suggesting that positive natural selection targets protein structure more strongly than protein sequence.

The success of the ENM in predicting the native dynamics of proteins solely based on the average structure of the native state (Tirion 1996, Atilgan et al. 2001) provides additional support to Str-CPE, since it suggests that the functional dynamics of proteins, which is an important factor for protein fitness, is largely determined by their native structure, underlying its importance for selection.

All these arguments concur in supporting Str-CPE. Nevertheless, so far no Str-CPE model can generate site-specific substitution models for phylogenetic inference, essentially due to the lack of an explicit model that predicts how any specific amino acid change at a given protein site modifies the native structure.

Here we develop such a Str-CPE model for predicting site-specific substitution processes. We find that this model is more restrictive than our previous Stab-CPE models, since it generates substitution processes with lower sequence entropy and lower substitution rates. In addition, the new Str-CPE model correctly predicts that completely exposed sites are more variable and evolve faster than partially exposed sites, and it provides higher likelihood to multiple sequence alignment (MSA) that include at least one protein of known structure. However, different from Stab-CPE models, the sequences represented by the stationary sequences of Str-CPE models on the average do not have a stable folded state.

By combining the Str-CPE model with our previous Stab-CPE model, we obtain an improved model characterized by lower tolerance to mutation, higher protein stability, higher likelihood of real MSA and higher folding stability. These findings suggest that the combined models represent selection on proteins more realistically than both models that ignore site-specificity and our previous Stab-CPE models. They also support the view that selection on protein structure conservation, which we represent through the parameter Λ_str_, is stronger than selection on protein folding stability Λ _stab_.

We refer collectively to our models as SSCPE (structure and stability constrained protein evolutioni) models. Next, we assessed the usefulness of the SSCPE models, finding that they infer more accurate phylogenetic trees than those inferred under the empirical substitution models, in the sense that the SSCPE-based trees are more similar to the phylogenetic trees of very distantly related proteins inferred with structural information.

Phylogenetic trees inferred under empirical models with among-sites substitution rate variation according to a gamma distribution (+G) sometimes show higher likelihood than the SSCPE models, in particular for alignments that contain more than 100 sequences. However, we show here that this higher likelihood is explained by their longer branches, which are positively correlated to the log likelihood score. This advantage disappears if we regularize the maximum likelihood (ML) fit of branch lengths, jointly optimizing the log likelihood and the sum of branch lengths, in the spirit of the minimum evolution principle and the maximum posterior probability principle. The assessment of the inferred trees presented here supports the view that this regularization, which we call regularized maximum likelihood and minimum evolution (REGMLAME), improves the results of phylogenetic fitting, as it is well-known for other types of regression.

We implemented the SSCPE models, and their application to phylogenetic tree reconstruction, into a computer framework named SSCPE.pl that is freely available from https://github.com/ugobas/SSCPE.

## Materials & Methods

### Model of structure change upon mutation

To model the deformation of the protein structure produced by a mutation, we adopt the linearly forced elastic network model (LfENM) proposed by Echave (2008). This model represents a mutation as a force *F*^mut^ that perturbs the native structure of the wild-type protein and it predicts the associated structural deformation vector Δ*r*_*i*_ as the linear response to this force of the elastic network model (ENM) that represents the equilibrium dynamics of the native state of the protein, i.e. the biologically active state, where the protein atoms perform small fluctuations around the average native structure represented in the PDB:

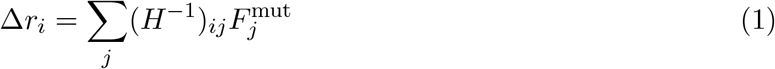

Here the indexes *i* and *j* label the 3*L* Cartesian coordinates of the C_*α*_ atoms that represent the *L* structured residues of the protein, *H*^−1^ is the inverse of the Hessian matrix of the second derivatives of the energy function (*H*^−1^ describes the dynamical correlations of the ENM) and *F*^mut^ is the force (perturbation) generated by the specific mutation.

Different from the original LfENM approach, which adopts an ENM that only moves the 3*L* Cartesian coordinates of the alpha carbons, here we adopt the torsional network model (TNM), an ENM in torsion angle space (Mendez and Bastolla 2010) implemented in the program tnm available at https://github.com/ugobas/tnm. The TNM moves all heavy atoms of the protein using as degrees of freedom a set of its torsion angles and constrains all other internal coordinates to their native values. In this work, we adopt the option that moves only the phi and psi backbone torsion angles, amounting to 2*L* degrees of freedom, fewer than the 3*L* ones of Cartesian ENM.

The TNM considers a contact-based energy function. All ENMs approximate the energy function as a quadratic function of the displacements, which is only valid when these displacements are small. In the TNM approach we add a quadratic potential that constraints the fluctuations of the torsion angles, 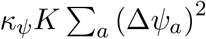, where *a* labels the torsion angles and *K* is an overall force constant. This additional potential allows to apply the TNM up to larger displacements representative of the thermal motion of the protein (Dehouck and Bastolla 2021). We adopt *κ*_*ψ*_ = 0.2, a value that maximizes the correlation between predicted atomic displacements and those observed in NMR ensembles (based on data from Dehouck and Mikhailov 2013).

The ENM provides an analytic description of the dynamics in the native state in terms of the normal modes, which are independent and collective harmonic motions of all protein atoms. Each normal mode *α* is characterized by its direction 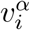 and by the frequency *ω*_*α*_ of the harmonic motions, which is inversely proportional to the amplitude of the thermal fluctuations of the normal mode (i.e. lower frequency normal modes contribute more strongly to thermal fluctuations). The number of normal modes equals the number of degrees of freedom excluding rigid body motions, i.e. 3*L* − 6 for the Cartesian ENM and 2*L* − 2 for the TNM. We express the structure change Eq.(1) in the base of the normal modes as

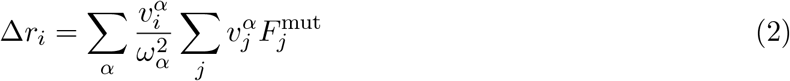

where *α* labels the normal mode, *j* labels the Cartesian components of all heavy atoms (not just alpha carbons), 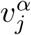 is the Cartesian component *j* of normal mode *α* (Mendez and Bastolla 2010), and we use the property of the normal modes 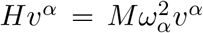. In the following, we denote by 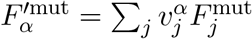 the projection of the mutation force on normal mode *α*.

### Representation of a mutation as a perturbing force

The LfENM models the force as the sum of components directed along the contacts that join the mutated residue *m* to its neighbours in the native state. These components are randomly drawn in the original LfENM approach. In contrast, here we model each component as the combined effect of the changes of amino acid size, contact stability and optimal contact distance produced by the mutation, in such a way that the mutation force depends explicitly on the amino-acids that are interchanged.

In the TNM a contact, designed as *C*_*ij*_ = 1, exists if the two closest heavy atoms belonging to residues *i* and *j*, indicated as *i*^′^ and *j*^′^, are closer than 4.5Å (this value was also determined by maximizing the correlations between predicted and observed atomic fluctuations). The vector joining the two residues is represented by the vector joining the two closest atoms, 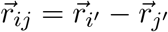.

We indicate each specific mutation at a protein site with the 3 indexes (*amb*), meaning that amino acid *a* (e.g. Leucine) at the mutated site *m* changes to amino acid *b* (e.g. Alanine). Each specific mutation generates a perturbation at each residue *k* in contact with the mutated site *m*, i.e. *C*_*mk*_ = 1. We indicate with *A*_*k*_ the amino acid at site *k* in the wild-type sequence. We consider three types of changes generated by these mutations: (a) Change of the amino acid size *s*(*a*) → *s*(*b*). (b) Change of the stability of the contact, which is expressed through the contact free energy matrix *U*, *U* (*a, A*_*k*_) → *U* (*b, A*_*k*_). (c) Change of the optimal equilibrium distance *d* between the amino acid pair, *d*(*a, A*_*k*_) → *d*(*b, A*_*k*_). Each of these changes generates the mutation force that is modelled as

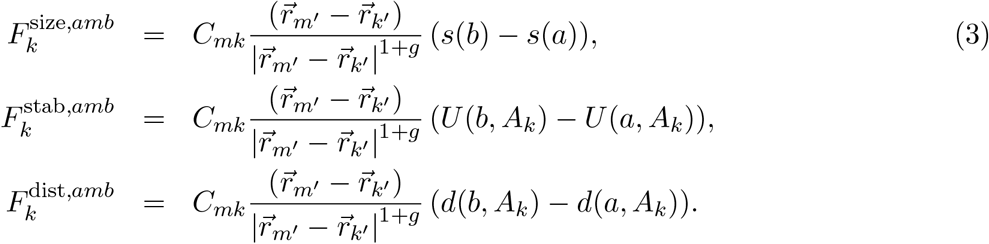

The components of the force are such that they tend to repel the two residues in contact if the mutated amino acid *b* is larger than the wild-type *a*, or the interaction energy *U* (*b, A*_*k*_) *> U* (*a, A*_*k*_) (meaning that the contact is destabilized), or the optimal distance increases, *d*(*b, A*_*k*_) *> d*(*a, A*_*k*_). These parameters are derived from protein statistics. *s*(*a*) is the number of atoms of each amino acid *a, U* (*a, b*) is the contact energy function derived in (Bastolla et al. 2000) and adopted for computing stability changes upon mutation in our previous Stab-CPE model (Bastolla et al. 2006, Arenas et al. 2015), and the optimal distances *d*(*a, b*) were determined as the most frequently observed distance in a non redundant subset of the PDB clustered at the 25% sequence identity level. They are available as Supplementary material. In this work we adopt the exponent *g* = 2, with the meaning that the mutation force is stronger at closer contacts. As in (Echave, 2008), the force at the mutated residue *m* is the reaction force of all of its contacts,

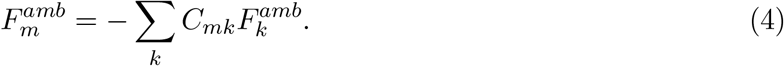

### Parameter optimization

For predicting the structural effect of mutations, we optimize three coefficients *W* that weight each type of contribution:

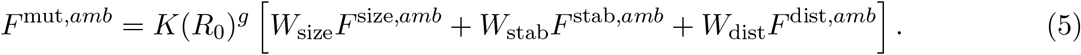

The multiplicative factor *K* (overall force constant) provides that the extent of the structural change does not vary when the force constant *K* is increased, which inversely decreases the fluctuations predicted by the TNM model. The factor (*R*_0_)^*g*^ with *R*_0_ = 3.25Å provides that the average extent of the structural change is almost independent of the exponent *g*.

The main difficulty for assessing and optimizing the mutation model is that the typical RMSD between the X-ray structures of the wild-type and of a mutant with only one amino acid change is of the same order as the RMSD between different X-ray structures with exactly the same sequence (Carpentier et al. 2019), and we do not know which part of the structural change is due to the mutation and which part is due to slightly different experimental conditions.

Here we take advantage of the fact that linear response theory also allows predicting the structural change in the absence of mutations. We recently proposed a null model of the random conformation changes produced by any change of experimental condition and that is not correlated with the native dynamics of the protein (Dos Santos et al. 2013). This null model is based on linear response theory, and it assumes that the contribution of each mode to the conformation change 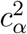 (i.e. the squared projection of the conformation change on normal mode *α*) is proportional to its contribution to thermal dynamics given by 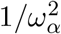. We verified this null model on a large set of proteins with same sequence and different structures (Dos Santos et al. 2013), finding that 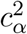 and 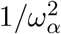 are indeed correlated for all pairs of conformations with identical sequence, but for functional conformational changes the low frequency normal modes contribute more than expected based on the null model, with the effect of reducing the harmonic energy barrier that opposes the conformation change (see also Dehouck and Bastolla 2021).

From Eq.(2), we see that the null model requires a perturbation *F*^NULL^ whose projection on each normal mode *α* is 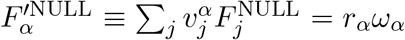, with *r*_*α*_ independent of the normal mode amplitude. Echave (2008) found that also the force *F*^mut^ that models the mutation has projections on normal modes that scale with *ω*_*α*_. This is so because the mutation is directed along the network of native contacts with a structure similar to that of the Hessian matrix *H* of the ENM. We may invoke a similar argument for justifying our null model. The final model for the observed mean square deviation (MSD) is

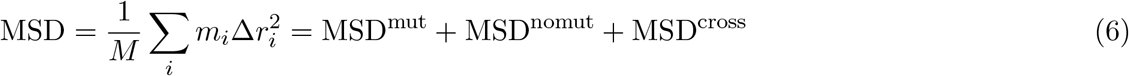

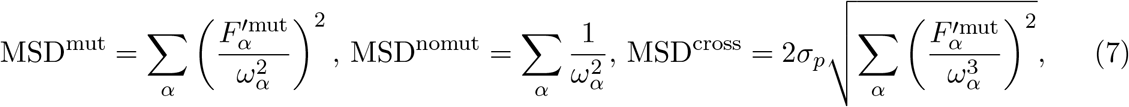

where *i* labels the Cartesian coordinates of the protein atoms, *m*_*i*_ is their mass, *M* is the total mass, *α* labels the normal modes, 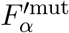 are the projections of the force on the normal modes, *σ*_*p*_ = ±1 is a binary random number, and we used the orthonormality conditions 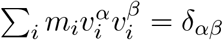. According to this model, the MSD of the mutant structure is given by three terms: (1) The MSD produced by the mutation force (mut), which depends on the mutation weights. (2) The null model fluctuations (nomut). (3) The cross term (cross) given by the scalar product of the deformations produced by the mutation and by the null model.

The cross term term cannot be neglected, and its sign cannot be predicted. We estimate it through its root mean square (RMS) value, Eq.(7) multiplied times the “random” sign *σ*_*p*_ = ±1 that is specific of each wild-type-mutant pair. We do not need to know *σ*_*p*_ for predicting the MSD due to the mutation, but only for assessing the prediction. To this end, we fit *σ*_*p*_ according to whether the observed MSD is higher or lower than the expected MSD based on the null model alone. Except for this sign, the MSD only depends on the mutation force and on the force constant *K*. We acknowledge that fitting *σ*_*p*_ may overestimate the accuracy of the model. For this reason, in this work we carefully evaluated the mutation model based on evolutionary data.

An additional problem is that sometimes the observed RMSD derives from functionally distinct conformations rather than from the effect of the mutation. Luckily, we can detect such instances and omit them from the training and test sets. Some pairs of structures significantly violate the null model of random conformational changes, since they present low frequency normal modes that contribute to the motion more than expected according to the null model. This implies that the harmonic free energy barrier that opposes to the motion is smaller than expected and the motion is more likely to happen spontaneously, suggesting that the observed conformation change is functionally important, such as an opening-close transition or ligand binding (Dos Santos et al. 2013, Bastolla and Dehouck 2019). Therefore, we eliminated conformational changes that present predicted harmonic energy barriers more than 50% lower than expected.

### Model of protein stability change upon mutation

This part of the model is the same as the one presented in (Arenas et al. 2015), and we recall it for completeness. We estimate the stability of the experimentally known native state of a protein adopting the contact matrix representation of its structure, *C*_*ij*_ = 1 if the residues *i* and *j* are in contact and zero otherwise, adopting the same definition of contact as in the TNM. Since contacts with |*i* − *j*| ≤ 2 are formed in almost all structures they do not contribute to the free energy difference between the native and the misfolded ensemble, and we set *C*_*ij*_ = 0 if |*i* − *j*| ≤ 2, while in the TNM model we set *C*_*ij*_ = 1 also for short range contacts.

The free energy of a protein in the mesoscopic state described by *C*_*ij*_ is modelled as the sum of contact interactions, *E*(*C,A*) = Σ_*i*<*j*_ *C*_*ij*_*U*(*A*_*i*_,*A*_*j*_), which depend on the types of amino acids in contact *A*_*i*_ and *A*_*j*_ and on 210 contact interaction parameters *U* (*a, b*). We use the parameters *U* (*a, b*) determined in (Bastolla et al. 2000).

For simplicity, we neglect the conformational entropy of the folded native state and estimate its free energy as 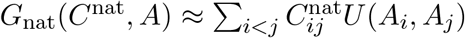. For the unfolded state, we neglect the contact interactions and estimate its free energy as *G*_*U*_ ≈ −*TLS*_*U*_, where *T* is the temperature in units in which *k*_*B*_ = 1, *L* is chain length and *S*_*U*_ is the conformational entropy per residue of an unfolded chain. We compute the free energy of the misfolded state from the partition function of the contact energy *E*(*C, A*) over a set of compact contact matrices *C* of *L* residues that are obtained from the PDB. In agreement with previous studies (Garel and Orland 1988, Shakhnovich and Gutin 1989), the resulting free energy is approximately described by the Random Energy Model (REM) (Derrida 1981),

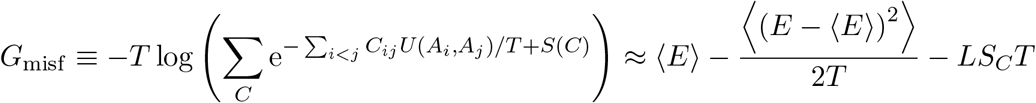

where *LS*_*C*_ is the logarithm of the number of compact contact matrices and ⟨.⟩ represents the average over the set of alternative compact contact matrices of *L* residues. We assume for simplicity that the conformational entropy, *S*(*C*_*ij*_), is approximately the same for all compact structures including the native one, and it can be neglected for computing free energy differences. This approximation is in good agreement with explicit computations of the free energy obtained through threading and real protein sequences present significantly lower stability of the misfolding ensemble than randomly reshuffled sequences (Minning et al. 2013). The previous estimate only holds above the freezing temperature of the REM (Derrida 1981), while the free energy remains constant below the freezing temperature.

Note that, although the free energy is defined in terms of Boltzmann averages, the result of the calculation depends only on the first two moments of the contact energies. We compute these moments from the corresponding moments of the contacts, which we pre-calculate when the program is initialized in order to accelerate the computations. This is important to achieve computational efficiency since the program must iteratively calculate averages over the site-specific distributions. In the initialization step, the program reads a file where we stored all the contact matrices of a non-redundant subset of the PDB, with only one representative for each cluster at 25% sequence identity, and it computes the contact statistics as described below. This step takes 45 sec for the longest protein with *L* = 1045.

The first moment of the energy, ⟨*E*⟩ =∑_*i<j*_ ⟨*C*_*ij*_⟩ *U* (*A*_*i*_, *A*_*j*_), depends on the moments of the contacts ⟨*C*_*ij*_⟩. To reduce the number of parameters, we assume that (*C*_*ij*_) depends only on the sequence distance |*i* − *j*| and on chain length *L*, ⟨*C*_*ij*_⟩ = *a*(*L*)*f* (|*i* − *j*|). We determine *f* (|*i* − *j*|) as the probability that two residues with sequence distance *i* − *j* are in contact, and the factor *a*(*L*) so that the average number of contacts for structures of length *L* is the same as for native structures (Minning et al. 2013).

For the contact correlation term, ∑_*ij*,*kl*_ (⟨*C*_*ij*_*C*_*kl*_⟩ − ⟨*C*_*ij*_⟩ ⟨*C*_*kl*_⟩) *U*_*ij*_*U*_*kl*_, we compute approximate averages over two sites and consider three kinds of terms.

1. 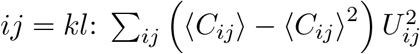 where we used that 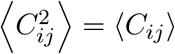.
2. We sum over *j* in contact *ij* and over *l* in contact *kl*: ∑_*ik*_ (⟨*n*_*i*_*n*_*k*_⟩ − ⟨*n*_*i*_⟩ ⟨*n*_*k*_⟩) *U*_*i*_ *U*_*k*_ where *n*_*i*_ = *∑*_*j*_ *C*_*ij*_ is the number of contacts of site *i* and *U*_*i*_ = ∑_*j*_ ⟨*C*_*ij*_⟩ *U*_*ij*_*/n*_*i*_ is its average energy.
3. We sum over *kl*: ∑_*ij*_ (⟨*C*_*ij*_*N*_*c*_⟩ − ⟨*C*_*ij*_⟩ ⟨*N*_*c*_⟩) *U*_*ij*_*U*, with *N*_*c*_ = ∑_*ij*_ *C*_*ij*_ total number of contacts and *U* = ∑_*ij*_ ⟨*C*_*ij*_⟩ *U*_*ij*_*/N*_*c*_ average energy of the whole chain.

Putting together these estimates, we obtain the free energy difference between the native and the non-native states as

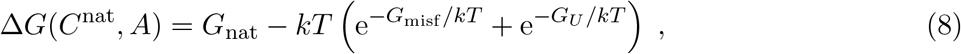

where the free energy of the non-native state is computed as a Boltzmann average, which is essentially equal to *G*_misf_ when the sequence is hydrophobic (*G*_misf_ − *G*_*U*_ */kT* ≪ −*kT*) and is essentially equal to *G*_*U*_ when the sequence is hydrophilic (*G*_misf_ − *G*_*U*_ */kT* ≫ *kT*). When we neglect stability against misfolding, we estimate Δ*G* = *G*_nat_(*C*^nat^, *A*) + *LS*_*U*_ as if all sequences were hydrophilic.

### Selection on protein structure and stability

The SSCPE models that we present here are based on the formal analogy between the Boltzmann distribution in statistical physics, where conformations are weighted by minus the exponential of the energy divided by *k*_*B*_*T* (i.e. *P* (*C*) = exp(−*E*(*C*)*/k*_*B*_*T*)*/Z*) and the stationary distribution in sequence space that arises from an evolutionary process with Fisher’s fixation probability. The latter distribution is proportional to the exponential of the logarithm of fitness *φ*(*A*) multiplied times a selection parameter Λ conceptually related with the effective population size *N*, *P* (*A*) = exp(Λ*φ*(*A*)) (Sella and Hirsch 2005, Halpern and Bruno 1999).

Similarly to the Boltzmann distribution, the sequence distribution can be formally defined as the distribution that maximizes sequence entropy with the constraint of given average value of the logarithm of fitness, ∑_*A*_ *P* (*A*)*φ*(*A*). This formulation is valid when all amino acids *a* have the same probability of being realized through the mutation process. Under a realistic mutation process, each amino acid will have a probability *P*^mut^(*a*) of being realized under mutation alone, and the condition of maximum sequence entropy must be generalized by the condition that the sequence distribution has minimal Kullback-Leibler divergence from the distribution under mutation alone, ∏ _*i*_ *P*^mut^(*Ai*).

Although the formulation in terms of a mutation process is elegant and is consistent with evolutionary theory, in practice it is difficult to cleanly separate selection at the amino acid level from mutation at the nucleotide level. The global (site-unspecific) amino acid frequencies reflect both mutational factors (number of codons and their stationary frequency) and unspecific selective factors (metabolic cost of the amino acids or their suitability to occurr in natural proteins) that are not taken into account by our model. To express this, we rename *P*^mut^(*a*) as *P*^glob^(*a*).

The above definition is only formal, because it is impossible to compute the normalization factor due to the huge number of possible sequences *A*. Nevertheless, this computation can be performed under the approximation that protein sites are independent (Arenas et al. 2015), which is almost necessary for computing the phylogenetic likelihood, and is commonly used in phylogenetic inference. If the fitness only depends on the properties of each individual protein site *i*, in other words if we can define a function *φ*_*i*_(*A*_*i*_) that evaluates the fitness of amino acid *A*_*i*_ at position *i* independent of all other amino acids, the stationary amino acid distribution may be represented as

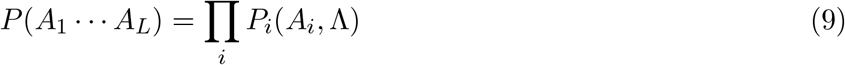

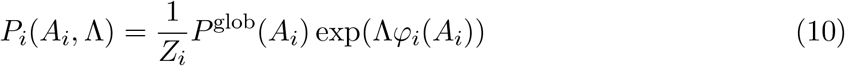

with *Z*_*i*_ = ∑_*a*_ *P*^glob^(*a*) exp(Λ*φ*_*i*_(*a*)).

### Stab-CPE mean-field model

In (Arenas et al. 2015) we introduced the mean-field (MF) model of stability-constrained protein evolution, in which fitness is proportional to the fraction of protein that is in the native state, which can be computed from the folding free energy of the protein Δ*G* (Goldstein 2011, Serohijos and Shakhnovich 2014, Bastolla et al. 2006) and the logarithm of the fitness is proportional to −Δ*G*. In the spirit of mean-field models of physics, the function *φ*_*i*_(*A*_*i*_) is determined self-consistently by averaging the exponential factor over the distributions generated by all other sites. In this way, we recover the influence of interactions between sites in a model where sites are formally independent:

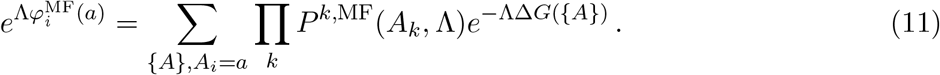

The MF distribution can be computed through an iterative algorithm whose complexity scales as the square of the number of sites and that converges rapidly, taking advantage of the precomputation of the statistics of misfolded conformations. We perform this computation with Λ = 1, obtain 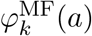 and then determine the optimal selection parameter Λ as explained in the next subsection.

### Stab-CPE wild-type model

Next, we define the stability constrained wild-type model (WT) that defines the fitness as minus the change in folding free energy (ΔΔ*G*) of the mutant protein 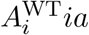 in which amino acid 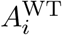 at site *i* of the wild type sequence is changed to *a*:

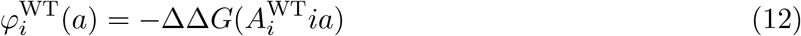

The WT model is mathematically simpler than the MF model, since it avoids the self-consistent calculation of the selection factors, and it produces results that agree well with the amino acid frequencies of extant protein families (Arenas et al. 2019). In fact, despite sequence space is exhaustively explored within the limit of the MF approximation, the native structure is maintained fixed, reducing the possible advantages of the more general MF approach.

### Str-CPE RMSD model

The structurally constrained models that we introduce here are similar in spirit to the WT model, since they are based on the predicted structural changes from the WT structure. In the RMSD model the logarithm of the fitness is modelled as minus the RMSD predicted by the tnm program for any specific mutation, i.e. the fitness decreases as the mutated structure becomes more different from the wild-type structure:

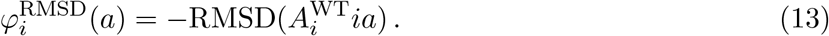

### Str-CPE DE model

We also evaluated the fitness function proposed by Echave (2008), whose logarithm is defined as minus the change in harmonic energy (DE) due to the mutation:

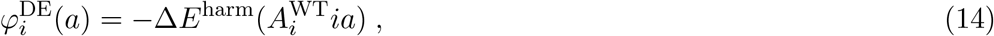

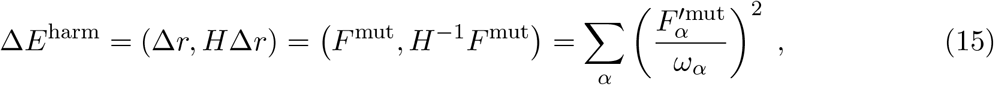

where *F*^mut^ is the perturbation generated by any specific mutation and *F*^′mut^ is its projection on normal mode *α*. The new version of the program tnm (Mendez and Bastolla 2010), computes and outputs the RMSD and the DE produced by all point mutations of the wild-type sequence if this computation is requested in the input file.

### Str-CPE for the wild-type residue

Note that the RMSD and DE deformations generated when the protein is mutated to the amino acid 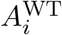 in the wild-type sequence in the PDB are zero. However, using this value as fitness would imply that the amino acid in the PDB has the maximal fitness under the structural point of view. Besides being unrealistic, this assumption produces very biased results under the point of view of phylogenetic inference that we inferred in this paper: The sequence represented in the PDB receives high likelihood, but the model infers phylogenetic trees that are very different from the reference tree, a clear sign of overfitting.

To address this problem, we compute it as the average deformation of the mutations to 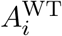 that start from all possible initial amino acids *a* at position *i*, assuming that they are distributed according to the stationary distribution *P*_*i*_(*a*) (with *P*^glob^(*a*) = 1*/*20 to avoid interferences with the estimate of this parameter) and that the deformation produced by the mutation 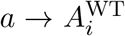 is the same as 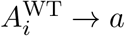:

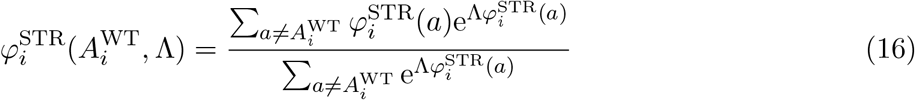

where STR represents either the RMSD or the DE model. Note that in this approach the fitness factors depend on the selection strength Λ.

### Combined Stab-CPE and Str-CPE models

Finally, we consider the four possible combinations of Stab-CPE and Str-CPE models, which we call SSCPE models:

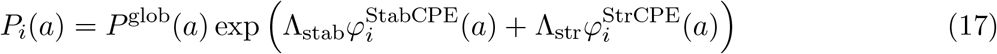

The SSCPE models have two selection parameters Λ_stab_ and Λ_str_.

### Cut-off probability

In phylogenetic applications it is important that the amino acid frequencies are not zero for any amino acid. Thus, we enforce the minimum frequency *P*_*i*_(*a*, Λ) ≥ *ϵ* = 0.001. The drawback is that with this condition the amino acid distribution is not an analytic function of Λ.

### Global frequencies

The SSCPE models take as parameters a vector of global amino acid frequencies *P*^glob^(*a*) and the selection parameter Λ. Given the latter, we determine the global frequencies for each SSCPE model and each value of Λ so that the frequency of each amino acid summed over all sites is equal to its frequency in the MSA:

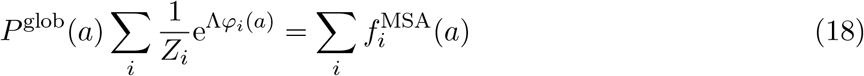

In this way, *P*^glob^ depends on Λ. Note that the normalization factor 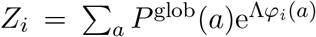 depends on *P*^glob^, so the equation must be solved iteratively. The above equation fits 19 parameters in such a way that the average frequencies are exactly equal in the model and in the MSA, and it is analogous to the +F option in phylogenetics.

### Selection parameter and regularization

The most important parameter of the site-specific substitution processes is the global selection parameter Λ. We explored two criteria for optimizing Λ:

1. Maximizing the log-likelihood of the site-specific amino acid frequency observed in the input MSA 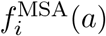 with respect to the model:

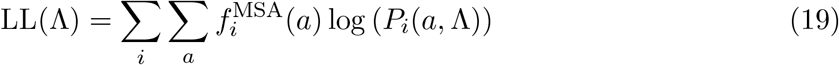
2. Minimizing the symmetric Kullback-Leibler divergence between the input MSA and the model:

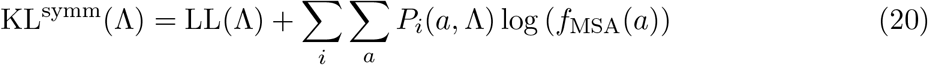

We apply a cut-off to the MSA so that its minimum value is 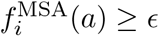.

We obtained better results with the KL criterion, which produces higher values of Λ and stationary frequencies that on the average represent more stable proteins. Therefore, we only present results obtained with this criterion.

Since the Str-CPE fitness 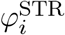 and the global amino acid frequencies *P*^glob^(*a*) depend on Λ, it is cumbersome to optimize the above scores analytically determining the value of Λ where the derivative vanish. We maximize them numerically through quadratic optimization, which is fast and accurate.

### Tykhonov regularization with specific heat criterion

As with all regression problems, it is important to regularize the fitted parameters in order to limit overfitting. In our experience, regularization is always beneficial since it reduces the cross-validation error of the fits and often eliminates non physical values, such as negative spring constants in the context of the ENM.

We apply the Tykhonov regularization, also called ridge regression (Hoerl and Kennard 1970), and we minimize the sum of the KL divergence plus a term that limits the value of the fitted selection parameter Λ, i.e. we minimize

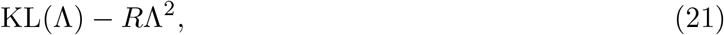

where *R* is the regularization parameter. Note that *R* is not a new fitted parameter, on the contrary it worsens the score achieved through the fit. Also note that this parameter is often denoted as Λ in the context of ridge regression, while here Λ is the parameter to be optimized. Given *R*, we determine Λ by minimizing the score either numerically or analytically, as the value at which the derivative vanishes.

In (Dehouck and Bastolla 2017) one of us noted the analogy between regularization and thermodynamics, in the sense that the error of the fit (here KL) can be interpreted as an energy, which increases when increasing the regularization parameter *R*, interpreted as a temperature. Therefore, the derivative of the error with respect to *R*, which can be shown to be positive, can be interpreted as a specific heat *C*_*V*_ :

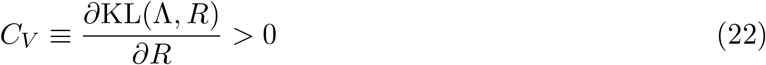

We found that, in many least square problems, the specific heat presents a maximum that separates a small *R* regime where overfitting is strong from a large *R* regime dominated by bias. The maximum of the specific heat presents good performances for different regression problems (Dehouck and Bastolla 2017, Bastolla and Dehouck 2019), and we adopt this criterion for determining the *R* parameter.Our program samples several values of *R*, computing the optimal value of Λ and the *C*_*V*_ numerically for each of them. Finally, it chooses the value of *R* with maximum *C*_*v*_ and refines its value through quadratic optimization.

### Site-specific sequence entropy and substitution rate

The sequence entropy at site *i* measures the variability of this site as

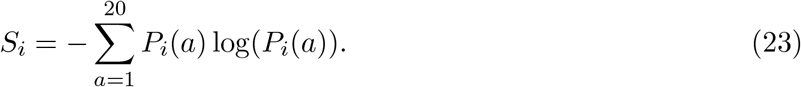

The site-specific substitution rates of the evolutionary models are computed as the weighted average of the substitution rate matrix *Q*_*i*_(*a, b*) = *E*_*i*_(*a, b*)*P*_*i*_(*b*),

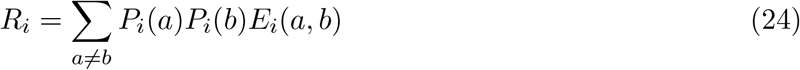

### Exchangeability matrix

Up to now, we did not describe how we define the exchangeability matrices *E*_*i*_(*a, b*) at each site *i* that are necessary for computing the site-specific substitution process *Q*_*i*_(*a, b*):

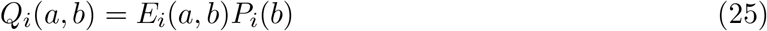

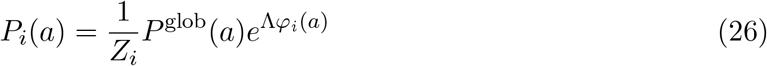

For computing the exchangeability matrices, we consider the following alternatives.

### Halpern-Bruno model

The simplest choice consists in choosing the same global exchangeability matrix for all the sites, *E*_*i*_(*a, b*) ≡ *E*^glob^(*a, b*). We call this option “noHB”. The alternative option, which we call “HB”, consists in applying the Halpern-Bruno model (Halpern and Bruno 1998) that expresses *E*_*i*_(*a, b*) as the product of a global substitution process *E*^glob^(*a, b*) that is the same at all sites times a site-specific fixation probability. To this end, following Halpern and Bruno we interpret *φ*_*i*_(*a*) as the logarithm of fitness and compute the fixation probability through the Fisher’s formula, which results in

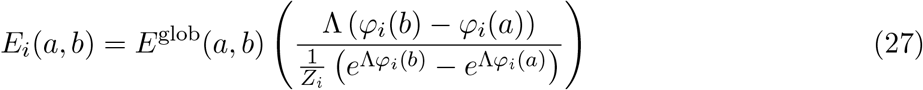

### Flux model

To determine the global exchangeability matrix 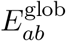, we also have two alternatives. The simplest one, which we call “noFL”, consists in choosing 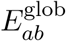 equal to one of the empirical exchangeability matrices such as JTT (Jones et al. 1992), LG (Le and Gascuel 2004) or WAG (Whelan and Goldman 2001): 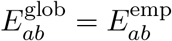.

The alternative option, which we call “FL”, consists in assuming that the empirical substitution matrices have to be trusted more in terms of the flux between any amino acid pairs that they determine, because this is the quantity that they fitted from empirical data. Thus, in the FL option we impose that the flux of any two amino acids *a* and *b* determined by the model, *R*_*i*_(*a, b*) = *P*_*i*_(*a*)*P*_*i*_(*b*)*E*_*i*_(*a, b*), and averaged over all sites *i* equals the flux measured in the empirical model for each pair of amino acids *a* and *b*, _*i*_ *R*_*i*_(*a, b*)*/L* = (*P*^emp^(*a*)*P*^emp^(*b*)*E*^emp^(*a, b*)). We get

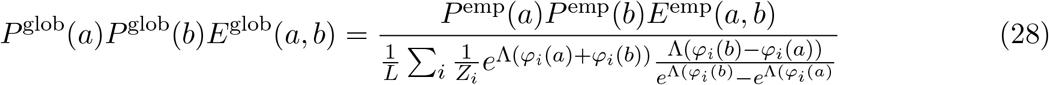

for the HB model, and

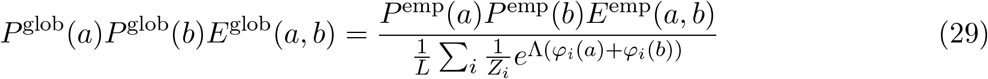

for the noHB model.

### Normalization of the substitution rate

Finally, another alternative arises from a technical feature of the program RAxML-NG, which internally normalizes the exchanggeability matrix at each site so that its substitution rate is always one: Σ_*ab*_ *P*_*i*_(*a*)*P*_*i*_(*b*)*E*_*i*_(*a, b*) ≡ 1. We explored two alternatives: either we allow the program to internally normalize the exchangeability (rate0) or we input to RAxML-NG a scale factor for each site so that, after the normalization, the exchangeability matrix is multiplied times this scaling factors that restores the substitution rate predicted by the SSCPE model (rate1).

As a result, we have eight different exchangeability models, corresponding to the combinations of HB/noHB, FL/noFL and rate1/rate0. We explored all these combinations, and we found the best results on the average with the options HB, FL and rate1, although for some score, model or data other combinations may yield better results. The presented results correspond to these options, unless otherwise state.

### Choice of the empirical substitution model

The empirical substitution model can either be chosen by the user or be internally optimized by the program, maximizing the following approximate likelihood function that does not consider the tree topology and assumes short branch lengths. The likelihood of the substitution between amino acid *a* and *b* in a pair of sequences that evolved through a time *t* is *P* (*a*) (e^*tQ*^)_*ab*_. At first order in *t*, this is approximately *tP* (*a*)*E*_*ab*_*P* (*b*) if *b* ≠ *a* and *P* (*a*)(1 + *tQ*_*a*,*a*_) otherwise, and its logarithm is approximately log(*t*)+log(*P* (*a*))+log(*P* (*a*))+log(*E*_*ab*_) or log(*P* (*a*))+*tQ*_*a*,*a*_. Summing over all pairs of aligned amino acids, whose number is *N* (*a, b*) and denoting *N* (*a*) the number of occurrences of amino acid *a*, we get:

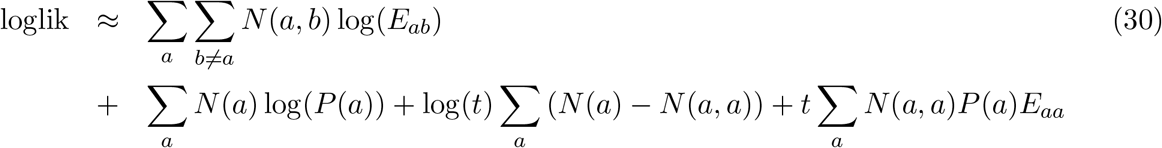

The value of *t* that maximizes Eq.(31) is readily determined where the derivative vanishes, *t* = − Σ_*a*_(*N* (*a*) − *N* (*a, a*))/Σ_*a*_ *N* (*a, a*)*P*_*a*_*E*_*aa*_ with *E*_*aa*_ = − Σ_*b*≠*a*_ *E*_*ab*_*P*(*b*). Plugging it into Eq.(31) we obtain the approximate value of the log likelihood for each of the three empirical substitution matrices implemented in our program (JTT, LG and WAG), from which we choose the optimal one.

### Assessment of the SSCPE exchangeability matrix

For given SSCPE model (8 types) and for the chosen empirical exchangeability matrix, the SSCPE model can adopt 8 combinations of 3 binary options of the site-specific exchangeability matrices (HB/noHB, FL/noFL, rate0/rate1). The program considers all of them and computes for each site the approximate likelihood given by Eq.(31). In this way, it assesses which combination provides the highest value of the approximate likelihood. This information is printed in the output file with extension SSCPE.summary.dat. The best combination is almost always HB+rate1, often in combination with FL and sometimes in combination with noFL. Through this score, we can also explore which SSCPE model presents the highest likelihood. However, not always the predicted model receives the highest phylogenetic likelihood in computations by RAxML-NG.

### Implementation

We implemented the computer program Prot evol that computes eight SSCPE models and prints their site-specific amino acid frequencies, summary statistics and site-specific substitution rates and sequence entropies.

We also implemented the framework SSCPE.pl that runs Prot evol or reads its results if they are present in the current folder and infers the maximum likelihood phylogenetic tree with the program RAxML-NG (Kozlov et al. 2019) for the required SSCPE model (possible choices: MF, WT, DE, RMSD, DEMF, DEWT, RMSDMF, RMSDWT, default: the model that obtains the best non-phylogenetic likelihood according to Prot evol) and the required options of the exchange-ability matrix (8 combinations of the binary options HB/noHB, FL/noFL, rate0/rate1, default: HB+FL+rate1).

We implemented the perl program K2 mat.pl, using as starting point the program K mat.pl (Soria-Carrasco et al. 2007), for computing the similarity between the inferred trees and a reference tree. Besides the K score based on one-parameter branch length fit and the topological score by Robinson and Foulds (1981) computed by K mat.pl, K2 mat.pl also computes the K2 score based on two-parameter branch length fit that is more appropriate for quantifying the similarity between sequence-based and structure-based trees (see Results).

Finally, we implemented the master script script run SSCPE.pl that, for an input MSA, runs a combination of several SSCPE models with several options of exchangeability matrix and several empirical substitution models with or without the options +F and +G. The script submits the computations to a queue system, which must be also modified by the user. After all computations are done, the master script must be called again with the option -examine and with a reference tree in Newick format (-tree) for comparing the trees inferred with each model to the reference tree with the program K2 mat. The results of these comparison, the log-likelihood computed by RAxML-NG, the sum of branch lengths and the REGMLAME scores are printed in a table called Results.[].txt.

Further details on the usage of the framework are presented in GitHub (see Data Avaibility).

### Concatenated alignments and multiple structures

The program Prot evol receives as input a protein MSA (-ali), and either an individual pdb file (-pdb) or a list of pdb codes (-pdblist) together with the folder where the pdb files are stored (-pdbdir). The corresponding structures may align to different regions of the MSA, which may be obtained by concatenating multiple proteins.

The program reads the MSA and the pdb files. Then it aligns each pdb sequence against each sequence of the MSA. If the best match has *<* 0.5 sequence identity the pdb structure is discarded, otherwise the program adopts the corresponding alignment ali and computes the folding free energy Δ*G*(*A*_*s*_, *C*_*p*_, ali) and the fitness *f*_*ps*_ = 1*/*(1 + exp(Δ*G*)) for each sequence *A*_*s*_ in the MSA aligned to the structure *C*_*p*_. Typically, the sequence of the pdb protein *s*(*p*) has Δ*G*(*A*_*s*(*p*)_, *C*_*p*_, ali) ≪ −1 and *f*_*ps*(*p*)_ ≈ 1. We discard the structures that are predicted to be unstable (Δ*G*(*A*_*s*(*p*)_, *C*_*p*_, ali) ≥ 0) or stabilize too few sequences (∑_*s*_ *f*_*ps*_ is lower than the minimum beltween 4.5 and 0.45 times the number of sequences in the MSA). Otherwise, the structure is used for computing the fitness factors 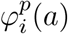 of the models MF, WT, DE and RMSD at the sites *i* of the MSA that are aligned with structure *p* and we compute the weight *w*_*p*_ ∝∑_*s*_ *f*_*sp*_ for obtaining the weighted average of 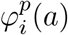 at sites with multiple structures.

Subsequently, we discard from the MSA the sequences for which the stability score ∑_*p*_ *f*_*sp*_ is less than 0.9, we remove the columns of the MSA that do not have any residue and relabel the columns of the MSA. Each such column *i* can either have a structural model and the corresponding fitness 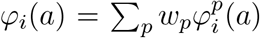 or it has *φ*_*i*_(*a*) ≡ 0 and it is described by the global amino acid frequencies *P*^glob^(*a*).

For each SSCPE model, the program considers several values of the regularization parameter *R*, optimizes the selection coefficients Λ numerically by minizing KL(Λ) + *R*Λ^2^, computes numerically the derivative *C*_*V*_ = *∂*KL*/∂*Λ, finds the value of *R* that maximizes *C*_*V*_ and recomputes the optimal Λ and the site-specific amino acid frequency 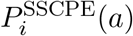, which it prints in a file called [].SSCPE.[MODEL].AA profiles.txt for the MSA columns that are structured, together with the global frequencies *P*^glob^(*a*) corresponding to that model.

Finally, for each model the program computes the site-specific exchangeability matrices according to the four models HB/noHB and FL/noFL, which depend on the site-specific frequencies, the global frequencies *P*_glob_(*a*) and the empirical substitution model, computes the corresponding approximate likelihood, and prints the four site-specific rates in a file called [].SSCPE.[MODEL].rate profile.dat. Since they can be recomputed rapidly, the exchangeability matrices are not printed to avoid to write large files. They are recomputed by SSCPE.pl for the required model and exchangeability options, then they are input to RAxML-NG as partitions and the correponding files are deleted upon completion of the phylogenetic computation.

## Results

### Predicting the amount of structure change caused by a mutation

We determined the weights *W*_size_, *W*_stab_ and *W*_dist_ in Eq.(5) by numerically fitting Eq.(7) over a training set of mutants with known structure. We omitted structures that are almost identical to the wild-type (WT), requiring that the root mean square deviation (RMSD) is at least 0.15Å. This condition is rather tolerant, but more restrictive conditions reduce the data set too much. From the starting set of 222 pairs, we eliminated 27 pairs of conformational changes that are likely to have a functional meaning (see Methods), obtaining 195 mutations with RMSD*>* 0.15Å and standard deviation 0.28Å. We randomly split the corresponding protein families into training set (156 pairs) and test set (39 pairs, i.e. 20%).

To avoid the influence of the binary parameter *σ*_*p*_, which we fit from the observed RMSD (see Methods), we optimized the correlation between the RMSD predicted only with the mutation and the observed RMSD, which is not influenced by *σ*_*p*_. Note that the value of *σ*_*p*_ does not influence the predicted RMSD produced by the mutation, which we need for modelling the substitution process. The parameters that result from the optimization are *W*_size_ = 3.05, *W*_dist_ = 15.6 and *W*_stab_ = 2.39, yielding the correlation coefficient between observed and predicted RMSD *r*_WT_ = 0.86 and root mean square error RMSE_WT_ = 0.20Å over the training set from wild-type to mutant, and *r*_mut_ = 0.75, RMSE_mut_ = 0.22Å from mutant to wild-type (the two sets differ since the torsional network model of the mutant is slightly different from the one of the wild-type). We evaluate the performances of the model on the test set, finding correlation coefficients *r*_WT_ = 0.95 and *r*_mut_ = 0.97 and RMSE_WT_ = 0.16Å and *RMSE*_mut_ = 0.11Å (see Supplementary Figure S1). These correlations were high because the test set contains only one pair with large RMSD, which is more difficult to predict. Considering only the mutation model we obtained poorer correlations (*r*_WT_ = 0.54, *r*_mut_ = 0.28 for the training set and *r*_WT_ = 0.68, *r*_WT_ = 0.65 for the test set), which supports that a combination of the mutation model, the null model and their cross product improves the prediction of the conformational change caused by a mutation. We will present more detailed results in a forthcoming publication.

For the sake of the evolutionary applications of the model, three general properties of the structural effect model, independent of the particular values of the parameters, are important. They are shared with the random force model proposed by Echave. The only advantage of the present model is that it can be explicitly applied to generate substitution models for phylogenetic inference.

1. Due to Eq.(3), the more contacts the mutated site *m* has, the more terms the resulting mutation force has. Therefore, mutating sites with more contacts tend to generate larger perturbations, and as a consequence these mutations are less tolerated. This is exemplified in Fig.1A, which shows the average RMSD generated by a mutation as a function of the number of contacts of the mutated site.
2. Due to Eq.(2), sites with more native contacts tend to be less deformed by the mutation because they produce lower flexibility in the ENM (their eigenvector components *v*^*α*^ are small in low frequency normal modes that contribute most to the thermal motion). As a consequence, the structural effect of mutation is weaker at highly connected sites, as observed in Fig.1B, which shows the RMSD of site *i* averaged over all possible mutations as a function of the number of contacts of the mutated site.
3. Finally, Eq.(3) implies that the RMSD caused by a particular mutation is correlated with the corresponding change of native contact energy, and the extent of this correlation grows with the parameter *W*_stab_.

**Figure 1:**
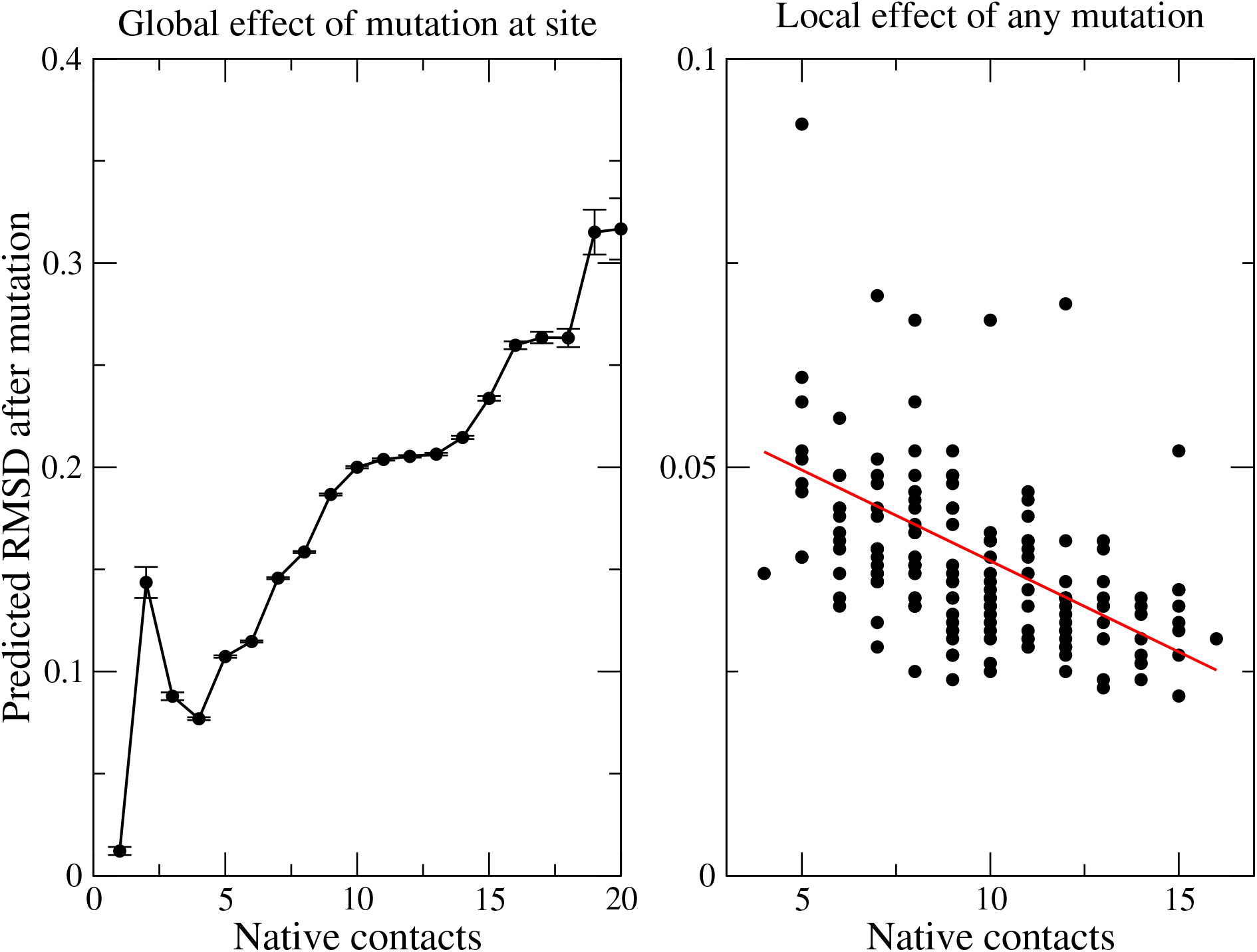
Effect of the mutation model as a function of the number of contacts of the mutated site (A) and the number of contacts at the position where the structure change is monitored (B)

### Assessment of the predicted amino acid frequencies

In this work, we compare our two previous Stab-CPE models MF (in which stability of amino acid *a* at site *i* is estimated self-consistently considering the amino acid frequencies at all other sites) and WT (in which stability is estimated considering the amino acids at all other sites in the wild-type sequence) against the new Str-CPE models RMSD and DE (in which we compute the RMSD and DE produced by amino acid *a* at site *i* adopting the ENM of the wild-type protein) and their four combinations RMSDMF, RMSDWT, DEMF and DEWT. They are summarized in table 1. We also consider the global (site-unspecific) amino acid frequencies determined by setting the selection parameters to zero (GLOB).

**Table 1:**
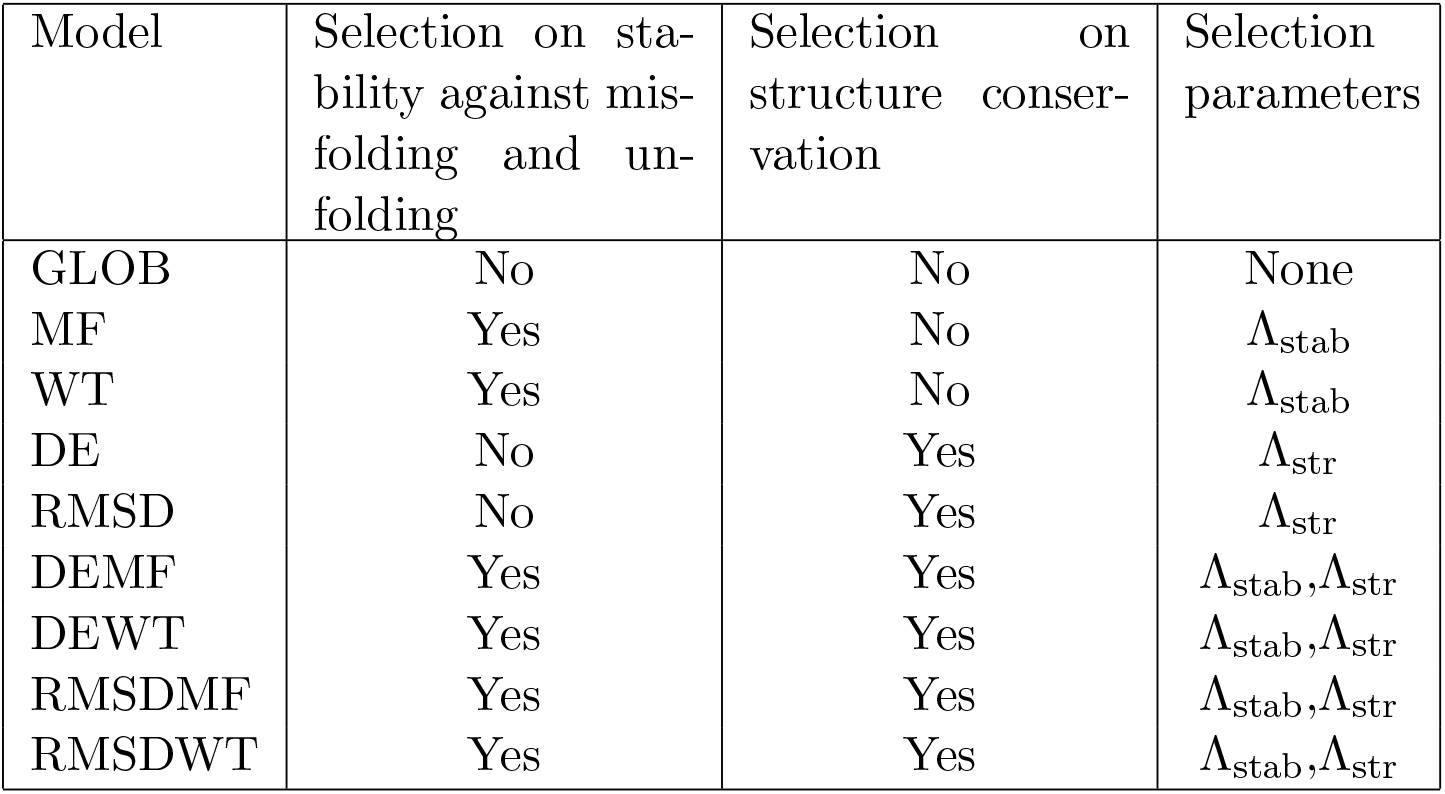
Definition of the SSCPE models studied in this work.

In Fig.2, we compare the results obtained with the site-specific amino acid frequencies computed with the 8 different SSCPE models on the MSA of 203 monomeric proteins. For all models, the global frequencies and the Λ parameter were determined minimizing the average of the symmetric Kullback-Leibler (KL) divergence between the amino acid frequencies of the model and the observed MSA at the regularization value determined with the “maximum specific heat criterion” (see Methods). However, setting a constant regularization value does not modify the qualitative results. We also consider the model GLOB obtained with the global amino acid frequencies setting Λ = 0.

**Figure 2:**
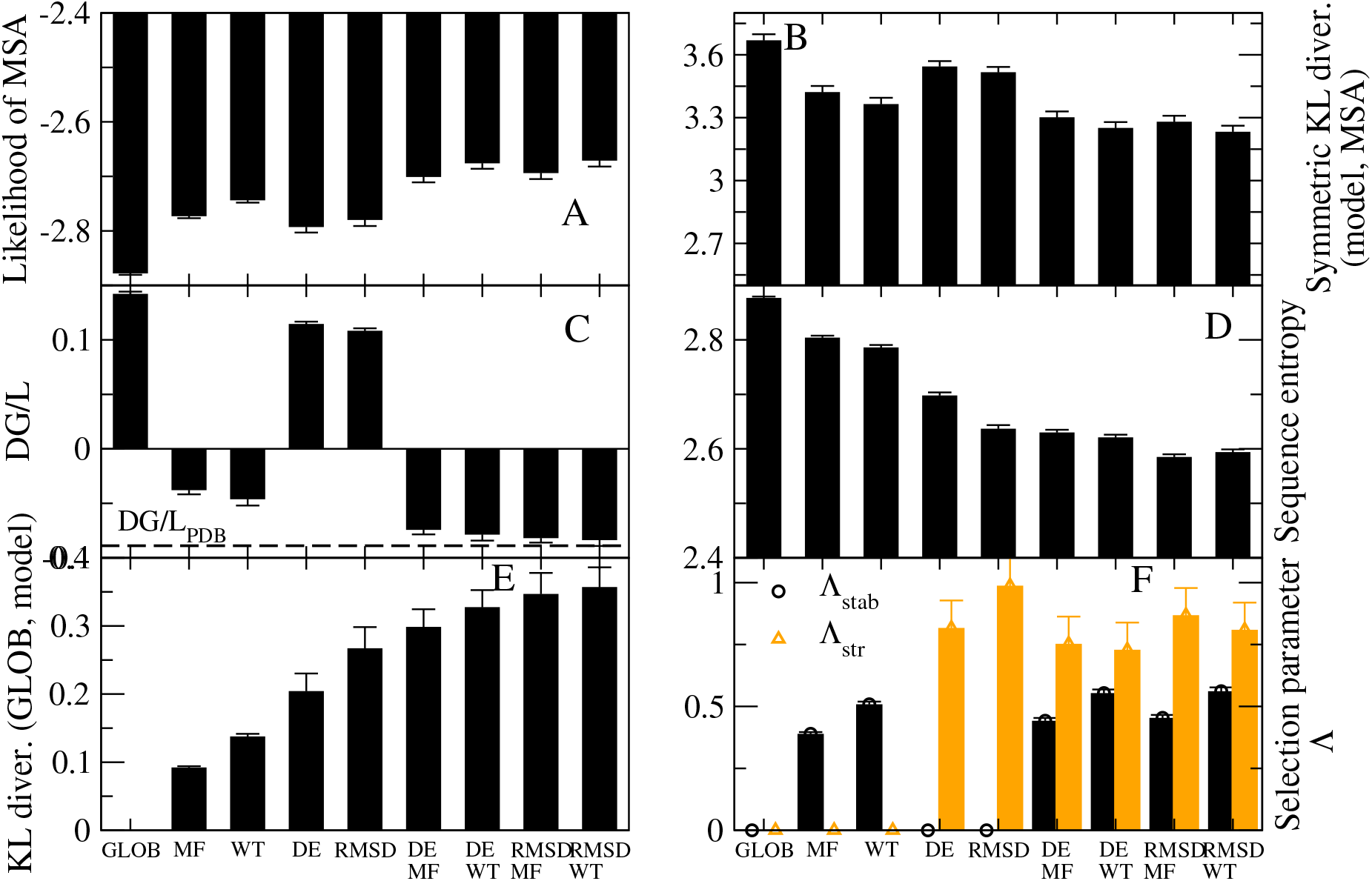
Assessment of the SSCPE profiles. Results obtained with eight SSCPE site-specific substitution models, listed in the horizontal axis, on a test set of 203 MSA with 10 to 300 sequences, at least one with known structure. A: likelihood of the observed MSA. B: Symmetric Kullback-Leibler divergence between each model and the observed MSA. C: Average folding free energy per residue of the model profile. D: Site-specific sequence entropy of the model profile. E: Kullback-Leibler divergence between the SSCPE model and the global model. F: Selection parameters. All shown quantities represent the average across all sites.

One can see from Fig.2A that the average likelihood of the MSA can be ranked as GLOB < DE *<* RMSD *<* MF *<* WT *<* DEMF ≈ RMSDMF *<* DEWT ≈ RMSDWT, i.e. Stab-CPE models generally present higher likelihood than the Str-CPE models and the combined models present higher likelihood, the highest being for the RMSDWT model. The same ranking is obtained with the KL divergence (Fig.2B), where the signs of the inequalities are inverted. Not surprisingly, Str-CPE models represent unstable proteins, since their average estimated folding free energy Δ*G* is positive, however the combined models are more stable than Stab-CPE models and their average Δ*G* is almost exactly the same as for the structures in the PDB (Fig.2C). Str-CPE models present lower entropy than Stab-CPE models (Fig.2D), which indicates that they are less tolerant to mutations, and the combined models show the lowest entropies. A similar message is obtained from Fig.2E, where one can see that Str-CPE models tend to deviate more than Stab-CPE models from the site-unspecific model GLOB, and from Fig.2F, where one can see that the selecion parameter associated to structure conservation (orange) tends to be larger than the one associated with folding stability (black). All these results suport the view that structure conservation is a stronger selective constraint than conservation of protein folding stability.

### Site-specific entropy and substitution rates

Fig.3 shows the average site-specific sequence entropy (A), substitution rate (B) and hydrophobicity(C) as a function of the number of native contacts for the different models and for real MSA, as reported by Jimenez-Santos et al. (2018). As previously observed by Jimenez-Santos et al. (2018b), the substitution rate is strongly influenced by the model of the exchangeability matrix. We tested the Halpern-Bruno exchangeability model (HB) versus a model without fixation probability (noHB), and the FLUX model in which, for each pairs of amino acids, their flux averaged across all sites is the same as the flux of the empirical substitution model versus the model noFL that adopts the exchangeability matrix of the empirical substitution model (Supplementary figure S2). The noHB option and the FLUX option tend to increase the rate at sites with many contacts, which are occupied by hydrophobic amino acids. In the main text, we show the substitution rates for the option HB noFL.

**Figure 3:**
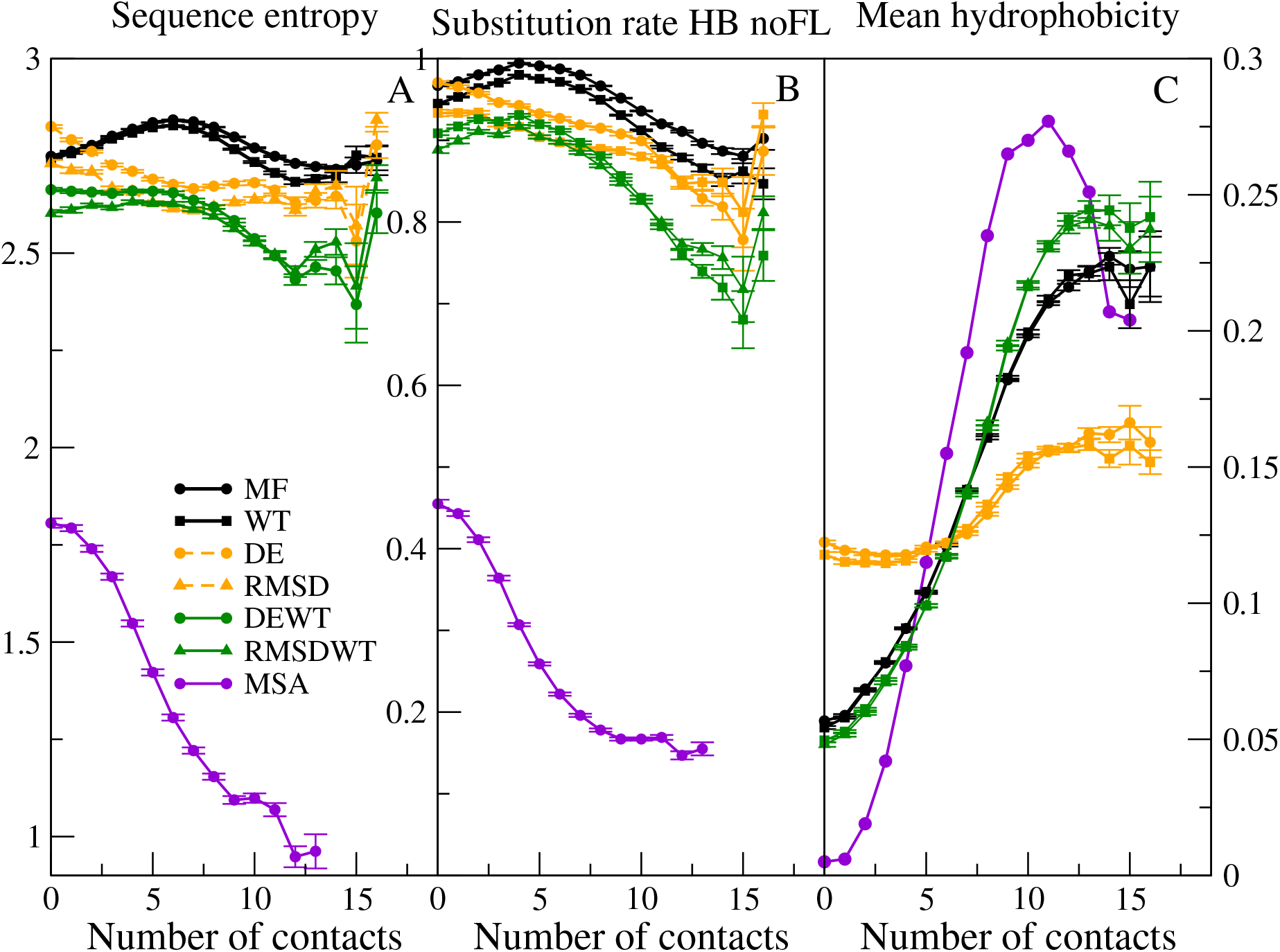
Sequence variability and hydrophobicity as a function of the number of contacts. Average site-specific sequence entropy (A), substitution rate (B) and hydrophobicity (C) as a function of the number of contacts of the site in the native state of the wild-type sequence (horizontal axis) for various SSCPE models. Data are averaged across 203 MSA with 10 to 300 sequences.

The entropy and the substitution rate of the Stab-CPE models MF and WT present a maximum for intermediate number of contacts (Fig.3A and B), indicating that these models are too tolerant for sites with intermediate number of contacts. In contrast, both sequence entropy and substitution rate tend to decrease with the number of contacts for the Str-CPE and SSCPE model except for sites with very high number of contacts, in qualitative agreement with observations from MSAs. Also the entropy and the substitution rate of the combined models present a maximum for intermediate number of contacts, although this maximum tends to occur before the one of Sta-CPE models. The combined models present lower entropies and substitution rates, i.e. they are less tolerant, although all models are quantitatively more tolerant than real MSA data.

Figure 3C represents the mean hydrophobicity of each site as a function of the number of contacts in the native state, for SSCPE models and for the real MSA. One can see that in all cases the hydrophobicity tends to increase with the number of contacts, as expected since this trend favors folding stability. This is even true for the Str-CPE models that do not select for folding stability, although in these cases hydrophobicity responds very weakly to the number of contacts. Interestingly the maximum hydrophobicity is attained before the maximum number of contacts (11 or 12 instead of 15) both for the MSA and for the RMSDWT model, which is the one that agrees best with the observed data. The hydrophobicity of the MSA spans a wider range than the models. In particular, it is lower than the models for small number of contacts and higher for high number of contacts, again suggesting that the models may be more tolerant than real protein evolution.

### Assessment of the SSCPE models for phylogenetic tree inference

We have shown that the new combined SSCPE models are more realistic than the site-unspecific model and our previous Stab-CPE models in terms of the dependence of site-specific evolutionary properties (sequence entropy, substitution rate, hydrophobicity) on the structural properties of the site, such as the number of native contacts, as well as in terms of the likelihod of the observed amino acids in the MSA and the average estimated folding free energy. We assess here the question whether these models are also useful for inferring more accurate phylogenetic trees. This is a difficult test, for which we cannot use simulations since, for distinguishing Stab-CPE and Str-CPE models we should simulate protein evolution with selection on the native structure, and we do not know any model that implements this. Therefore, we decide to compare the trees inferred with SSCPE and empirical substitution models to the trees derived with structural information with the program PC_ali (Bastolla et al. 2023). This program adopts an integrated measure of protein similarity in sequence (sequence identity) and structure (TM score, i.e. fraction of superimposed residues (Zhang and Scolnick 2004) and contact overlap, which does not require spatial superimposition of the structures), uses it for constructing a hybrid multiple alignment, and uses the corresponding evolutionary divergences for constructing a Neighbor-Joining tree (NJ, Saitou and Nei 1987). Note that PC_ali does not adopt the models of folding stability and structural changes on which the SSCPE model is based, and it infers phylogenetic trees through the NJ algorithm while we infer trees with SSCPE adopting the ML approach. Therefore, approximating the structure-based trees with sequence-based trees is a difficult test.

We determined PC_ali trees for five subsets of distantly related proteins, related at the super-family level, whose properties are summarized in Table 2. The reference phylogenetic trees are iluustrated in Supplementary Figures S3-S7. One can see that they are highly consistent with the functional classification of proteins in terms of Gene Ontology and Enzyme Commission numbers, which supports their good quality.

**Table 2:** Properties of the five studied superfamilies. Number of sequences, mean branch length of the structure-based NJ tree and ME tree, correlation between log-likelihood and K2 score from the NJ tree (negative means that higher likelihood trees are more similar to the reference), correlation between regularized likelihood and K2 score from the NJ tree (positive means that trees with better regularized score are more similar to the reference), correlation between log-likelihood and K2 score from the ME tree, correlation between regularized likelihood and K2 score from the ME tree.

| Superfamily | Sequences | Mean<br>branch<br>length<br>NJ | Mean<br>branch<br>length<br>ME | Corr<br>(loglik,<br>K2(NJ)) | Corr<br>(reglik,<br>K2(NJ)) | Corr<br>(loglik,<br>K2(ME)) | Corr<br>(reglik,<br>K2(ME)) |
| --- | --- | --- | --- | --- | --- | --- | --- |
| Aldolase | 21 | 0.327 | 0.195 | -0.91 | 0.87 | 0.33 | -0.31 |
| Globins | 64 | 0.137 | 0.072 | -0.61 | 0.54 | -0.39 | 0.46 |
| NADP | 84 | 0.246 | 0.131 | -0.22 | 0.19 | 0.53 | -0.30 |
| Ploop_C1 | 48 | 0.210 | 0.117 | -0.08 | -0.05 | 0.49 | 0.00 |
| Ploop_C2 | 14 | 0.339 | 0.199 | -0.43 | 0.49 | 0.31 | -0.27 |

We compared the reference NJ trees determined through structural information with the ML trees determined with 12 empirical models: JTT (Jones et al. 1992); LG (Le and Gascuel 2008); WAG (Whelan and Goldman 2001), with all four combinations of the options +F and +G) and the eight SSCPE models with the options HB, FLUX and rate (we also studied other options, but present here only these ones for the sake of simplicity because they produced on the average better results). We adopted three metrics: (1) The normalized topological distance (RF; Robinson and Foulds 1981). (2) The normalized K-score dissimilarity (Soria-Carrasco et al. 2007), which assumes that the branch lengths of the comparison tree are given by the corresponding branches of the reference tree multiplied times a fitted scaling factor. The K score consists of the mean square error of the fit. With respect to the RF distance, it has the advantage that it penalizes little the discrepancies that occur at short branches. Since protein structure is known to diverge more slowly than protein sequence (Illergard et al. 2009, Pascual-Garcia et al. 2010), we expect that the scaling factor of the K score is less than one. (3) Previous work suggested that protein structures diverge very slowly when the protein function is conserved, but they diverge faster when protein function changes, because of positive selection that targets the protein structure (Pascual-Garcia et al. 2019). Therefore, we extended the K score to the K2 score that adopts two scaling factors, which are interpreted as taking care of branches where protein function is conserved and where protein function changes. For this sake, we developed the perl program K2 mat, using as starting point the program K mat (Soria-Carrasco et al. 2007).

The results are presented in Fig.4. Fig.4A shows that the average K score is in general lower for the SSCPE models (black bars) than for the empirical models (orange bars). This difference is more accentuated when we examine the K2 score (Fig.4B), whose value is much smaller than the K score, supporting its better quality. Indeed, in the figures we rank the models according to their K2 score. The two-parameter fit is supported by at least two of the criteria AIC, BIC or corrected AIC for almost all models and superfamilies (Supplementary Figure S8). Interestingly, the best model in terms of average K2 score is the SSCPE model that maximizes the likelihood for each superfamily (ML(SSCPE)), followed by the simple Stab-CPE model MF and then the combined models RMSDMF, DEMF, DEWT. The Str-CPE models RMSD and DE score the worst among the SSCPE models, but still they perform better than the empirical models. The empirical model showing the maximum likelihood for each superfamily (ML(emp)) is one of the worst in terms of tree distance, which seems in contradiction with the good score of the ML(SSCPE) models.

**Figure 4:**
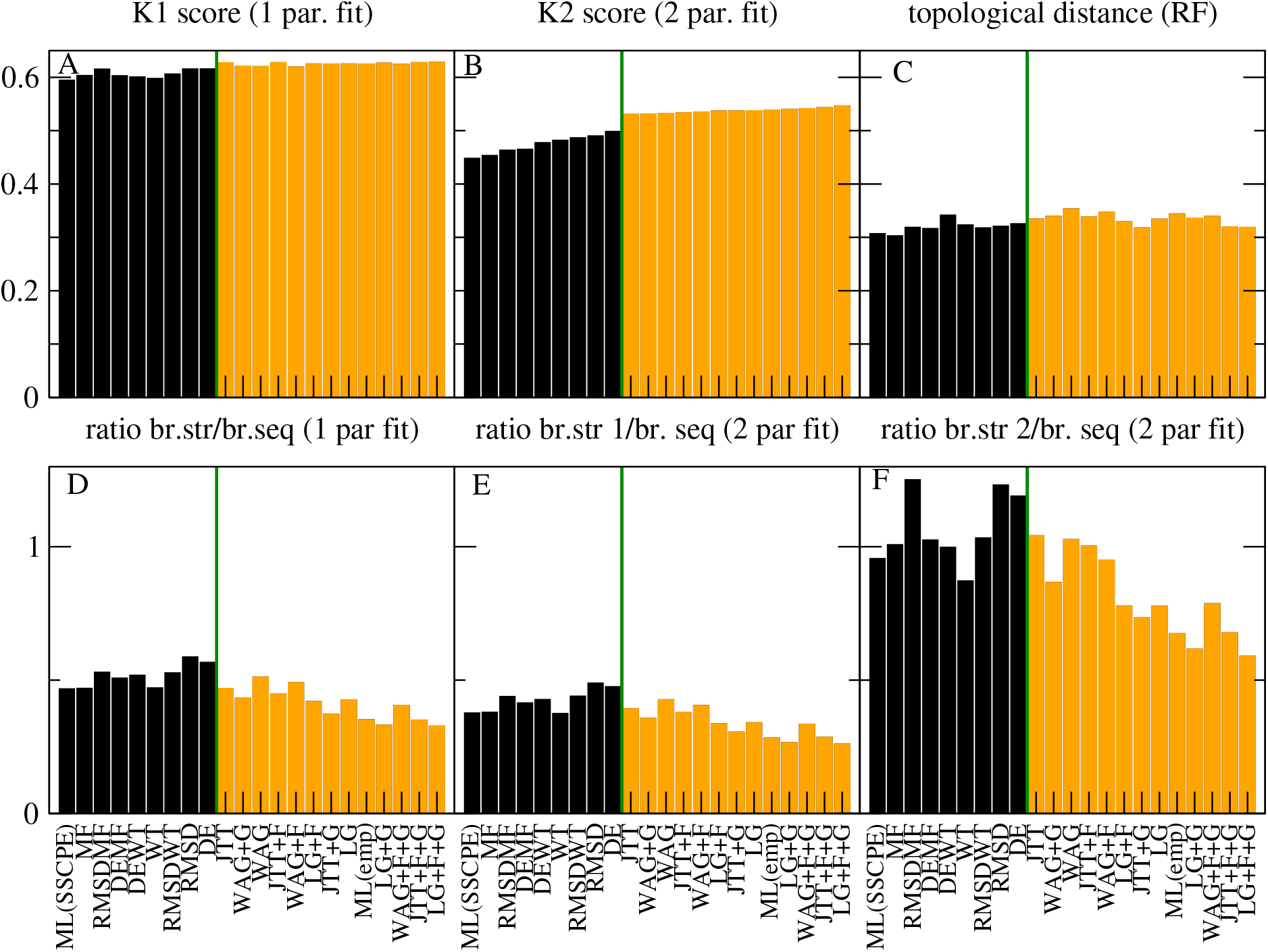
Assessment of SSCPE and empirical models with NJ structure-based phylogenetic trees. Comparisons between the NJ structure-based trees of five superfamilies and the corresponding ML trees obtained with SSCPE models (black bars) or empirical models (orange bars). A: Normalized K score, B: Normalized K2 score, C: Normalized topological distance (Robinson-Foulds) between structure based and sequence-based branches. Fitted ratios between the structure-based and sequence-based branches for 1 parameter fit (D), two parameter fit and short structure branches (E) or long structure branches (F). The models are ranked by smaller K2 scores. Each value is averaged over five superfamilies. The SSCPE models are run with the options HB, FLUX and rate all active.

As expected, the scaling factor of the one-parameter fit, which represents the ratio between structure divergence and sequence divergence, is less than one for all models and all superfamilies (Fig.4D). The two-parameter fit evidences slow branches in which the scaling factor is even smaller (Fig.4E) and fast branches in which the scaling factor approaches one (Fig.4F). Inspection of the phylogenetic trees supports the view that the latter branches represent events in which the protein function, represented by its GO term or by the EC numbers, changed. We will present a more quantitative analysis of this important property in another publication.

### Regularized maximal likelihood and minimum evolution

We then tested whether the presented results are robust when, instead of the NJ trees adopted by our program PC_ali, we use minimum evolution (ME) trees inferred with the program FastME (Lefort et al. 2015) applied to the structure-based divergence matrices generated by PC_ali. By visual inspection of the trees (Supplementary Fig.S3-S7) we realized that in some cases the GO terms or EC numbers support the NJ tree rather than the ME tree, despite it is frequently claimed that ME trees are more accurate (Lefort et al. 2015). Therefore, we sought an objective score for selecting between the two types of tree.

In the ML inference methods, the branch lengths *t*_*b*_ are obtained through a fit. It is well known that fits are threatened by overfitting, and their performances (for instance in terms of cross-validation) improve considerably when they are regularized by penalizing large values of the fitted parameters, for instance through the Tykhonov regularization that we adopt here for regularizing the fit of the selection parameter Λ. In the context of phylogenetics, the parameters are the branch lengths and, by the mathematical definition of the likelihood function, there is a clear trade-off that longer branches tend to be associated with higher likelihood. In this case, the L1 regularization that constrains the absolute values of the parameter is equivalent to minimizing the following quantity:

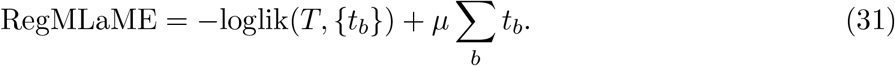

In other words, the regularized log likelihood also integrates the minimal evolution principle that is adopted for instance by the FastME program. Furthermore, the regularized log likelihood score can be seen also as the maximum posterior probability (MPP) of a tree *T* with an exponential prior distribution of branch lengths, while the ML tree can be seen as the MPP tree with uniform prior distribution, which seems to be less justified than the exponential distribution. Therefore, the regularized ML and ME tree unifies the three criteria most frequently adopted for tree inference: ML, ME and Bayesian. Finally, in (Dehouck and Bastolla 2017) one of us noted the analogy between regularization and thermodynamics. In the present context, −*log*.*lik* can be interpreted as an energy, which increases when increasing the regularization parameter *µ*, interpreted as a temperature, so that the quantity represented in Eq.(31) can be interpreted as a free energy.

If we fit branch lengths with ML for different trees using the same model, we find that they are positively related with the sum of branch lengths, loglik ≈ *µ ∑*_*b*_ *t*_*b*_ + *c*. This fitted parameter *µ* represents a natural choice for the regularization parameter *µ*. This choice amounts at not rewarding the likelihood in Eq.(31) unless it exceeds what is expected given the branch lengths of the tree.

For ME fits, the situation is the opposite. They are biased to favour short trees, but the tradeoff is that they present low likelihood. Also in this case, Eq.(31) provides a less biased score than the length of the tree. We adopted this score for comparing the NJ and ME trees. We proceeded as follows: first, we computed the likelihood of the trees with the model WAG+F+G, which is the ML model for most superfamilies. In this way, we verified that the ME trees have much lower likelihood than the NJ trees, as expected. Second, we recomputed the branch lengths maximizing the likelihood while maintaining the topology of the tree. In this way, we obtained four trees with their branch lengths and likelihood scores that are positively correlated, as expected (see Fig.5A to E, squares). Third, we computed the parameter *µ* for each superfamily from the regularized fit of the log likelihood versus the branch lengths and, with this value, we computed Eq.(31) for all trees, choosing the one with minimum value. For three superfamilies (Globins, NADP, Ploop C1) this is the ME tree, but for the remaining two it is the NJ tree. We did not modify its branch lengths, since they contain important structural information.

**Figure 5:**
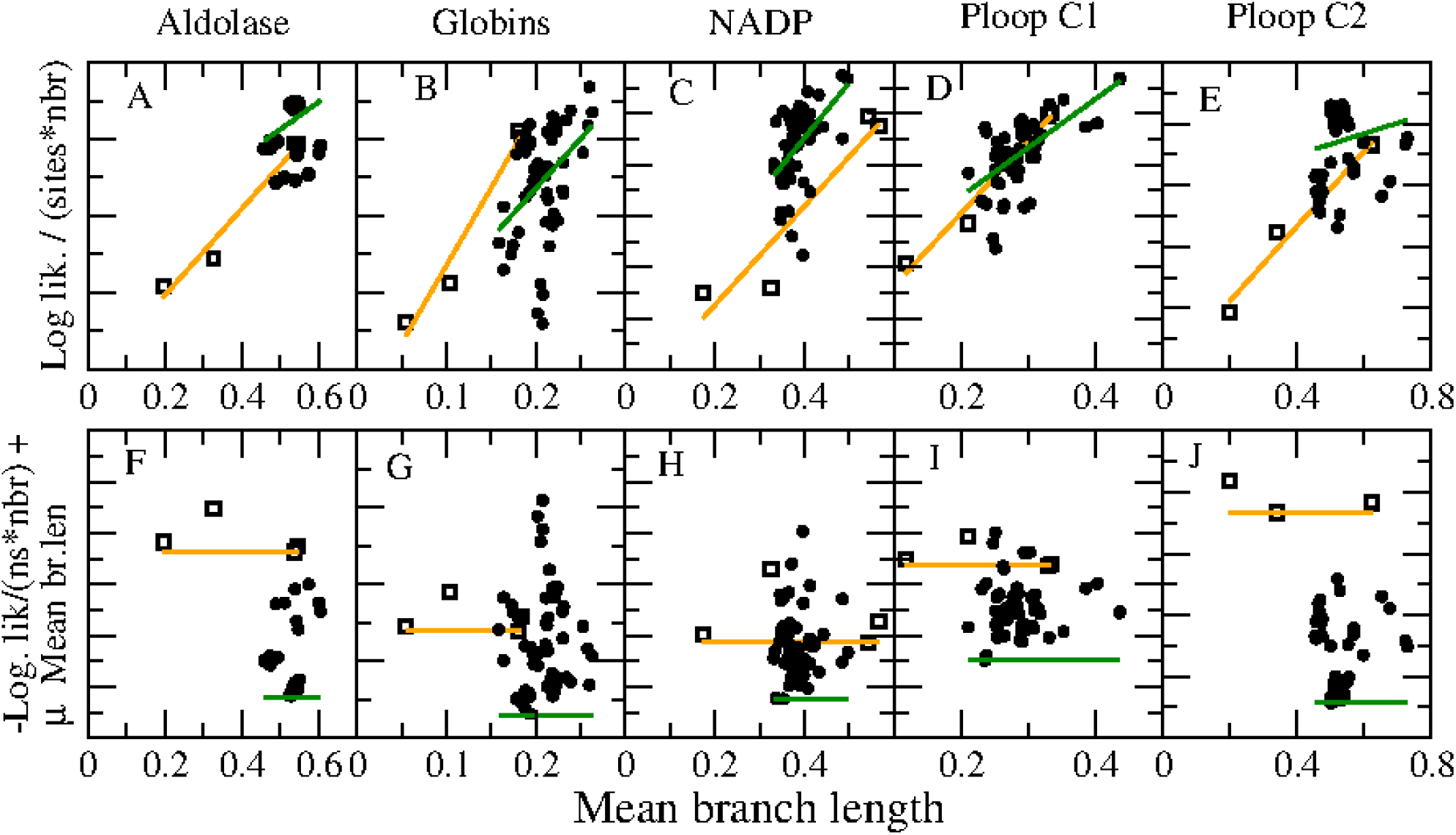
Selection of the reference phylogenetic tree. A-E: For the neighbor joining (NJ) and minimal evolution (ME) structure-based trees, we compute the log-likehood adopting the WAG+F+G model and recompute the branch lengths through maximum likelihood, obtaining four trees (squares) that evidence the trade-off between branch length and log-likelihood. A similar relationship holds for log-likelihoods of the ML trees computed with each SSCPE and empirical model (back symbols) versus their branch lengths. The fits of log-likelihood versus branch lengths (orange lines) estimates the regularization parameter *µ* with which we compute the REGMLAME score (figures F-J). For each superfamily, we choose as reference the structure-based tree with minimum REGMLAME score.

We then compared the ML trees inferred with the empirical models and with the SSCPE models to the reference trees obtained with the best between the NJ and ME approach. Despite the reference tree changed in three out of five cases, the qualitative results are robust and they generally improve, since all the dissimilarities K, K2 and RF decreased (compare Fig.6 with Fig.4). The K2 scores are better supported with the new reference trees for the SSCPE models (see Supplementary Figure S8B), although they are slightly worse supported for the empirical models. We also analyzed the relationships between the dissimilarity scores and the log likelihood and the REGMLAME scores of all ML trees built with different models (see Table 2 and Supplementary Figures S9 and S10). For NJ trees, as expected the log likelihood scores are negatively correlated and the REGMLAME scores are positively correlated with the dissimilarity scores for all superfamilies except Ploop C1 for which the correlations are not significant. The absolute value of the correlations is larger for the log likelihood score than for the regularized score, probably because the latter was not computed in a consistent manner (the branch lengths were not obtained by minimizing Eq.(31). However, for the ME trees the correlations have the opposite sign, positive between dissimilarity and log likelihood and negative between dissimilarity and Eq.(31). This supports the view that the branch lengths in ME trees are strongly biased.

**Figure 6:**
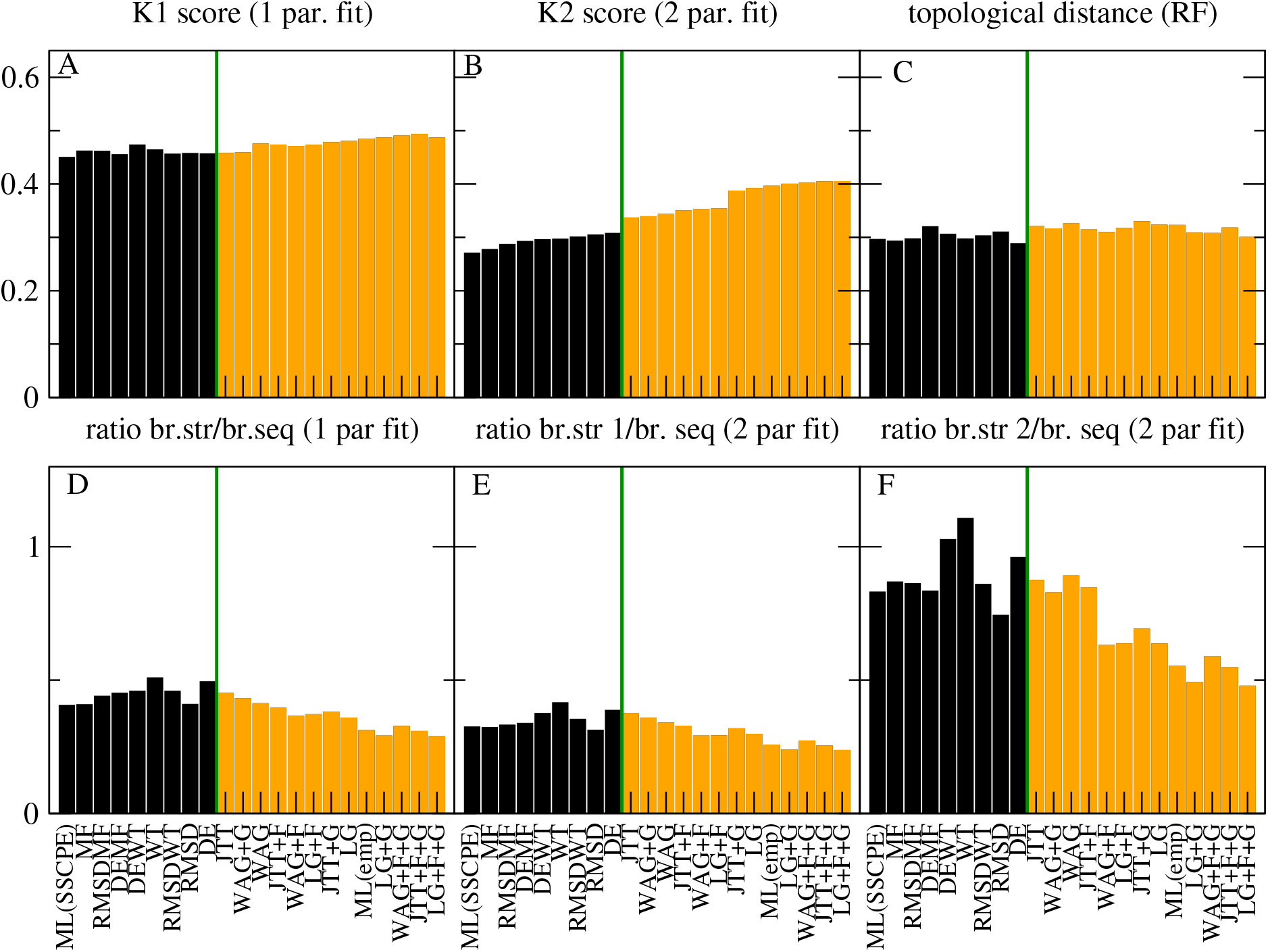
Assessment of SSCPE and empirical models with selected structure-based phylogenetic trees. Comparisons between five reference structural trees (best between NJ and ME tree) and the corresponding ML trees obtained with SSCPE models (black bars) or empirical models (orange bars). A: Normalized K score, B: Normalized K2 score, C: Normalized topological distance (Robinson-Foulds) between structure based and sequence-based branches. Fitted ratios between the structure-based and sequence-based branches for 1 parameter fit (D), two parameter fit and short structure branches (E) or long structure branches (F). The models are ranked by smaller K2 scores. Each value is averaged over five superfamilies. The SSCPE models are run with the options HB, FLUX and rate all active.

### Comparison of the phylogenetic likelihood of the models

We compared the empirical models and the SSCPE models under the point of view of the likelihood scores. Fig.7B shows that the average likelihood of the SSCPE models (black bars) is always larger than the one of the empirical models (orange bars) without the +G option, but it is smaller than some of the empirical models with the +G option. However, this higher likelihood of the +G empirical models is attained at the price of longer branch lengths (Fig.7C) and when this is taken into account the SSCPE models (except DE and RMSD) always have lower REGMLAME score than the empirical models (Fig.7A. In all figures, the models are ranked according to the REGMLAME score).

**Figure 7:**
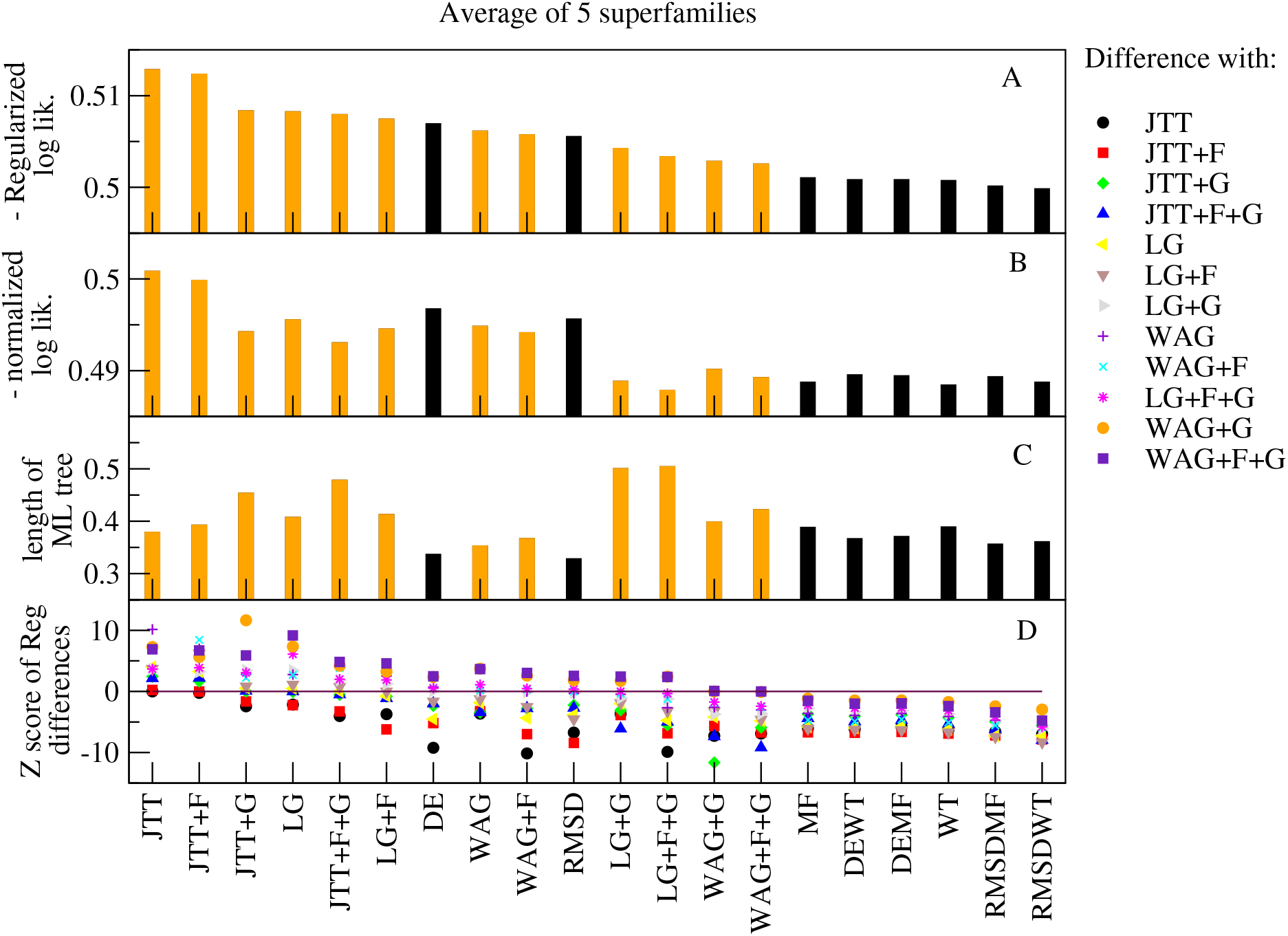
Regularized likelihood scores of SSCPE and empirical models for the super-family MSAs. Substitution models, listed in the horizontal axis, are ranked according to their regularized log-likelihood (REGMLAME) score averaged over the five superfamilies. Empirical substitution models are orange, SSCPE substitution models are black. A: REGMLAME score. B: Minus normalized log-likelihood. C: Average branch length of ML trees. One can see that low REGMLAME scores trade-off high likelihood and short branch length. D: Z score of the difference of the REGMLAME score between the named model and each of the empirical substitution models.

We caution that, in this and the following section, the computation of the REGMLAME score is only approximate because the branch lengths should be computed minimizing Eq.(31) and the parameter *µ* should be fitted separately for each model. However, these results are consistent with the finding that the SSCPE model perform better than empirical models, as assessed through the comparisons with the structure-based trees.

Finally, we used the site-specific substitution processes to infer phylogenetic trees of 10 randomly chosen protein alignments of our database (Supplementary Table S1) with the ML method implemented in the program RAxML-NG (Kozlov et al. 2019). The results are presented in Fig.8. In this case all of the +G empirical models present higher likelihood than the SSCPE models (Fig.8B), which is particularly so for large trees with more than 100 sequences (Supplementary Fig.S12B and D). The increased likelihood of the option +G is partly compensated by the biased increase of the branch length (Fig.8C), so that the REGMLAME score is more uniform (Fig.8A). The WAG+G model has still lower regularized score than any SSCPE model for large alignments (100 residues or more), except for the smallest one (1iu4, 10 sequences) that presents an advantage for SSCPE models so large that the minimum value of the average REGMLAME score is attained by the models RMSDWT and RMSDMF when we consider 1iu4. We removed this special case from Fig.8 in order to avoid to bias the results. Again, we note that the REGMLAME score that we compute here is only approximate.

**Figure 8:**
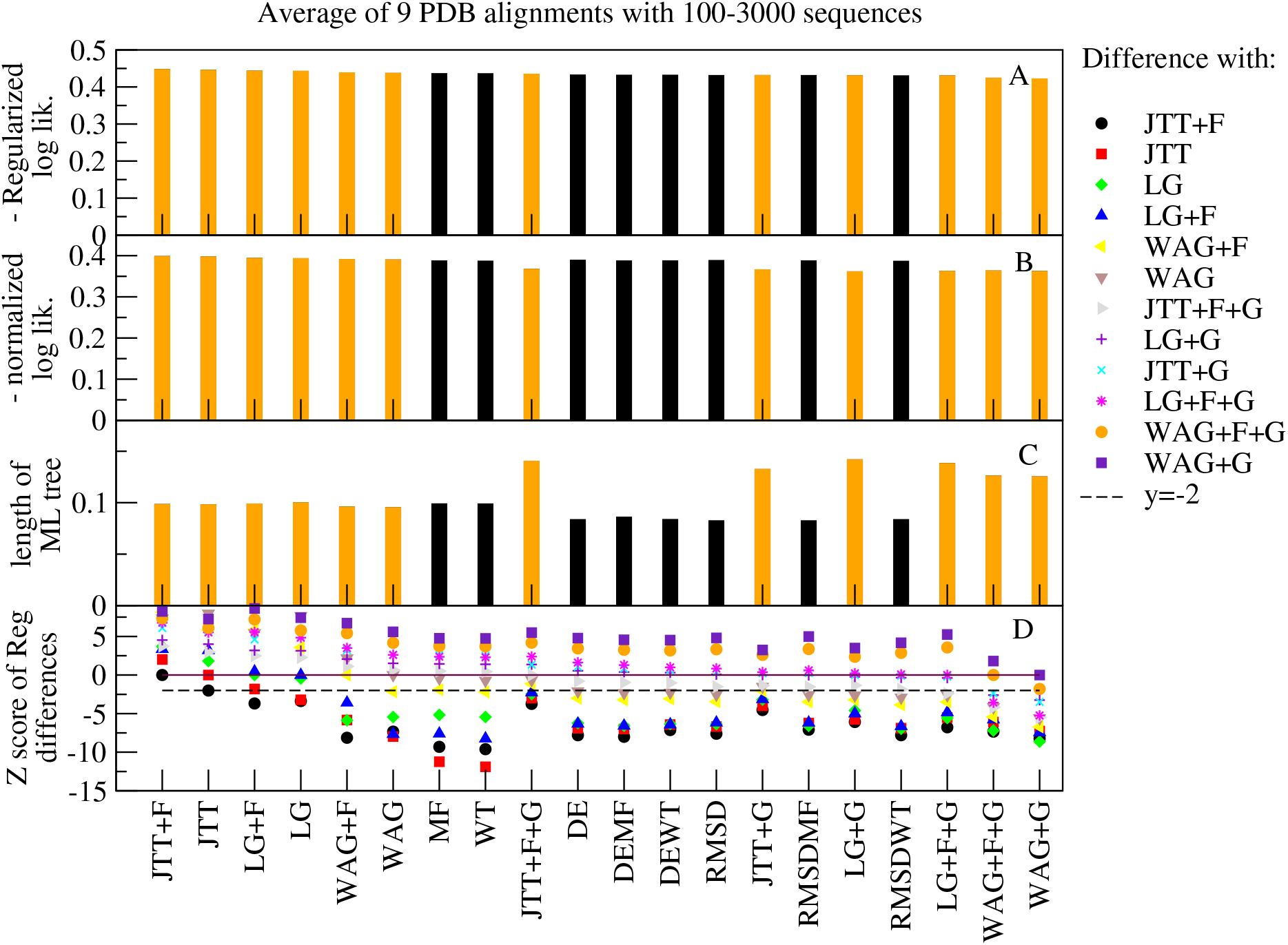
Regularized likelihood scores of SSCPE and empirical models for 9 MSAs with known native structure. We excluded 1iu4 because in this small MSA (10 very similar proteins) the empirical models score much worse than the SSCPE models, biasing the averages. Substitution models, listed in the horizontal axis, are ranked according to their regularized log-likelihood (REGMLAME) score averaged over nine multiple alignments of proteins with known structure. Empirical substitution models are shown in orange, SSCPE substitution models are shown in black. A: REGMLAME score. B: Minus normalized log-likelihood. C: Average branch length of ML tree. One can see that low REGMLAME scores trade-off high likelihood and short branch length. D: Z score of the difference of the regularized log-likelihood between the named model and each of the empirical substitution models.

## Discussion

Although the selective forces that act in protein evolution are complex and difficult to model, many years of modelling and observations have revealed general biophysical principles that most natural proteins must obey (Liberles et al. 2012, Wilke 2012, Sikosek and Chan 2014). However, the empirical substitution processes that are commonly used for phylogenetic inference ignore these biophysical principles, and they fit the site-specific variability of substitution rates through the gamma model (Yang, 1993), which is unaware of the protein structure. Here we show that introducing biophysical realism in the substitution models of protein evolution can improve phylogenetic inferences.

### Stability-constrained versus structure-constrained protein evolution

The folding stability of the native state is a biophysical property that received much attention in the literature (see for review Goldstein 2011, Serohijos and Shakhnovich 2014, Bastolla et al. 2017). Previous work produced site-specific stability-constrained substitution models that can be used for phylogenetic inference (Arenas et al. 2015). However, these Stab-CPE models assume for simplicity that the native structure is strictly conserved in evolution, and they cannot represent the selective process for maintaining it.

The native structure strongly influences native protein dynamics (Tirion 1996, Atilgan et al. 2001) and it is thought to be under strong negative selection when the protein function is conserved (Illergard et al. 2009, Pascual-Garcia et al. 2010) and under strong positive positive selection when the protein function changes (Pascual-Garcia et al. 2019). Therefore, it is crucial to consider it for modelling natural selection. In this work, we introduced two structurally constrained fitness functions based on the model by Echave (2008) that predicts the structural changes due to mutations as the linear response of the elastic network model that represents the protein dynamics to the perturbation caused by the mutation. Advancing with respect to Echave’s proposal, we developed an explicit model of the specific amino-acid mutations in terms of changes of size, stability and optimal distance of the native contacts formed by the mutated residue. Irrespective of the parameters, and similar to Echave’s original model, our model predicts that mutations at sites with many contacts induce larger deformations (Fig.1A) and sites with many contacts tend to respond less to perturbations (Fig.1B).

We introduce here two Str-CPE models that represent the logarithm of the fitness as minus the predicted RMSD due to the mutation (RMSD model) or as the predicted change in the harmonic energy of the elastic network (DE model, similar to the original proposal by Echave). We compared these models with two variants of our previous Stab-CPE models that model the logarithm of the fitness as minus the folding free energy Δ*G* evaluated with contact interactions, and consider as non-native states both the unfolded state and compact but incorrectly folded states (Minning et al. 2013).

These structure and stability constrained (SSCPE) evolutionary models depend on the global amino acid frequencies (as in empirical models with +F), which we determine imposing that the mean amino acid frequencies across all sites equal the global amino acid frequencies, and on the selection parameter Λ. We fit Λ by minimizing the symmetric Kullback-Leibler (KL) divergence between the site-specific amino acid frequencies of the MSA and the equilibrium frequencies of the models. To reduce overfitting, which favour large values of Λ and attributes very small probabilities to amino acids that are not observed, we regularized the fit by the Tykhonov regularization scheme (Hoerl and Kennard 1970), also known as ridge regression, choosing the regularization parameter *R* that maximizes the derivative of the score with respect to *R* (Dehouck and Bastolla 2017), inspired by the analogy between regularization problems and statistical physics.

The resulting site-specific SSCPE models provide lower KL divergence, lower sequence entropy and more stable proteins than the global model with optimized frequencies and Λ = 0 (Fig.2A-D). We found that Str-CPE models are characterized by lower entropy, diverge more from the site-unspecific global model and tend to have larger selection parameter Λ than the Stab-CPE models (Fig.2D-F). These results support the view that selection on protein structure is stronger than selection on protein stability. Nevertheless, the sequences described by the Str-CPE models are not stable on the average (Fig.2C) and they describe worse the amino acid distributions in the MSA (Fig.2A-B). Next, we combined the Str-CPE and Stab-CPE models. These combined models present better agreement with empirical MSAs, lower entropy, higher folding stability and diverge more from the site-unspecific model than both Stab-CPE and Str-CPE models (Fig.2A-E), showing consistent improvement under all criteria.

We examined site-specific evolutionary properties as a function of the number of contacts of the site in the native structure. While Stab-CPE models present maximum tolerance to mutations in terms of both sequence entropy and substitution rate at intermediate number of contacts, which qualitatively disagrees with observations (Jimenez-Santos et al. 2018), the Str-CPE and SSCPE models predict that the tolerance to mutations tends to decrease with the number of contacts and agree better with observed data (Fig.3A, B). Moreover, SSCPE models produce a more realistic range of mean hydrophobicity (Fig.3C). Nevertheless, from the quantitative point of view, all these substitution models of protein evolution appear to be more tolerant to mutations than the MSAs that we examined, which present lower sequence entropies and substitution rates. This may indicate that the models still do not capture important selective constraints, but it may be also in part attributed to insufficient sampling of the protein sequences, which underestimates the site-specific variability.

In conclusion, the combined SSCPE models represent site-specific selection on protein sequences more realistically than Stab-CPE and Str-CPE models, and much more than the global model that does not consider site-specific selection.

### Phylogenetic inference

We then tested whether the SSCPE models improve phylogenetic inference. We obtained the structure-based reference trees of five data sets of proteins distantly related at the superfamily level with the program PC_ali (Bastolla et al. 2023), and we inferred the maximum likelihood (ML) trees of the same proteins with several combinations of SSCPE and empirical substitution models with the program RAxML-NG (Kozlov et al. 2019).

We compared the structure-based and the sequence-based trees by fitting the lengths of the corresponding branches either with only one scale parameter (K score, Soria-Carrasco et al. 2007) or with two scale parameters (K2 score, our own extension of the Soria-Carrasco et al. program). For all models and all superfamilies, we found that the branches of the structure-based trees tend to be shorter than for sequence-based trees, in agreement with the notion that protein structures tend to evolve more slowly than sequences, but some of the branches are almost equally long in structure and in sequence, in agreement with the notion that structure evolution accelerates when the protein function changes (Pascual-Garcia et al. 2019). This feature may provide a mean of detecting changes of protein function, which we plan to explore in the future.

We found that the site-specific SSCPE models tend to infer trees that are more similar to the reference trees in terms of the K score, in terms of the topological distance by Robinson and Foulds (1981), and above all in terms of the K2 score, which is more relevant in this context since it takes into account the acceleration of protein structures upon function change.

### Regularized maximum likelihood and minimal evolution (REGMLAME) score

We first considered structure-based trees inferred with the neighbor joining method (NJ, Saitou and Nei 1987), and later tested whether these results are robust under structure-based trees inferred with the minimal evolution method (ME, Lefort et al. 2015). However, by direct inspection of the ME trees, it was apparent that sometimes their quality was not as good as those of the NJ trees. We devised a function for scoring them in an objective way.

For doing so, we used the program RAxML-NG for computing the likelihood of the structure-based trees under the WAG+G empirical model. We observed that the log likelihood is strongly posiively correlated with the sum of branch lengths, as expected from the mathematical definition of the likelihood. In fact, the likelihood is computed from the elements of the matrix e^*tQ*^, where *t* is the branch length, and for different amino acids the logarithm of the likelihood is positively related with *t*. Thus, there is a trade-off between log likelihood and branch length. A natural way to determine the optimal balance is by regularizing the fit of branch lengths, penalizing large values of the parameters (i.e. long branches) or, equivalently, penalizing short branches with low likelihood. This regularization leads to minimize the REGMLAME score, Eq.(31). It is well known that regularizing the fitted parameters improves their ability to generalize and reduces the occurrence of unphysical values (see for instance (Hoerl and Kennard 1970, Dehouck and Bastolla 2017, Bastolla and Dehouck 2019)), and we expect that this holds true also for the regressions used for phylogenetic inference. The REGMLAME score can also be interpreted as a maximum posterior probability under a prior distribution of trees that is exponential of the branch lengths. Therefore, the REGMLAME score unifies the three most used tree inference principles: ML, ME and MPP.

Applying an approximate version of the REGMLAME score, we found that the NJ trees have better score for two superfamilies, and the ME trees have better score for the remaining three. When we compared the SSCPE and empirical trees to the selected reference trees, we found that the comparison scores consistently improved and once again the phylogenetic trees inferred under the SSCPE models were more similar to the reference phylogenetic trees than the phylogenetic trees inferred under the empirical substitution models.

Finally, we computed the likelihood scores and the branch lengths of the ML trees obtained with 12 empirical substitution models and 8 SSCPE models. Empirical models with the option +G sometimes present higher likelihood than the SSCPE models. However, this improvement is achieved at the expense of biased longer branches. When this is taken into account with the REGMLAME score, the SSCPE models score better.

### Software implementation

Since the computations of the site-specific substitution matrices are fast and the site-specific matrices can be readily handled by the programs RAxML-NG or PAML, these results support the use of these structure-aware and selection-aware models for phylogenetic inferences in the many cases in which a representative structure is available in the PDB or can be predicted by AlphaFold2 (Jumper et al. 2021). The program tnm is the computational bottleneck, because of the computation of the normal modes and the 2 × 19*L* deformations produced by all possible mutants, which scales as *L*^3^. The tnm results are stored and reused if needed. The computer time is 9.7 sec for a small protein with *L* = 129 amino acids (PDB 132l), 51.9 sec for an intermediate one of 217 residues (1zio) 153.9 sec for one with 314 residues (1hqc) and 3598 sec for a large protein with 907 amino acids (1vlb). The program Prot evol computes the 8 SSCPE models in 7.1 sec for 132l, 9.3 sec for 1zio, 11.7 for 1hqc and 109.9 sec for 1vlb. To reduce the computation time, users may run only Stab-CPE models for large proteins and Str-CPE and combined models for small ones.

## Data Availability

The framework SSCPE.pl that performs phylogenetic tree inference under the SSCPE models with the program RAxML-NG is freely available at https://github.com/ugobas/SSCPE. Given an alignment of protein sequences and a list of representative protein structures (i.e., PDB files), SSCPE.pl uses the output of Prot evol or runs the program if the output does not exist, and performs phylogenetic tree inference with RAxML-NG under the specified SSCPE model.

SSCPE.pl can be installed by downloading the package SSCPE.zip at https://github.com/ugobas/SSCPE/blob/main/SSCPE.zip that provides all the needed codes and compiles and installs the programs tnm, Prot evol and SSCPE.pl. Instructions for the installation and the use are present at https://github.com/ugobas/SSCPE. The program RAxML-NG is included for completeness, but we recommend to install the latest version from https://github.com/amkozlov/raxml-ng.

The package SSCPE.zip also contains the program K2 mat, which compares two phylogenetic trees by performing two-parameter fit of the corresponding branch lengths under a modification of the program K mat.pl (Soria-Carrasco et al. 2007) and the master script script run SSCPE.pl that, for given MSA, runs a combination of several SSCPE models and empirical substitution models, submits the computations to a computing cluster, and compares the resulting trees with a reference tree computing several other scores, including an approximated REGMLAME score.

Supplementary figures and data are stored in the Dryad data repository at http://datadryad.org. They include: 1. Supplementary Figures S1-S12 described in the main text. 2. Mutation parameters used for tnm predictions. 3. MSAs of the 203 studied proteins. 4. RMSD and 5. DE predicted by the tnm program for all possible mutations of the studied proteins. 6. Data generated for testing the models: Site-specific substitution rates, sequence entropy, mean hydrophobicity and number of contacts. 7. Summary data for all tested models and all proteins. 8. For each superfamily, structure-based phylogenetic trees and ML trees inferred with the different empirical models and SSCPE models. Table with the comparisons between the reference structure-based trees and the ML trees of all models. 9. For each of the 10 protein families studied for phylogenetic inference, ML trees inferred with the different empirical models and SSCPE models. Table with the likelihood, sum of branch length and regularized likelihood of the ML trees of all models.

## Funding

This work was supported by the grants PID2019-109041GB-C22/10.13039/501100011033 and PID2019-107931GA-I00/AEI/10.13039/501100011033 of the Spanish Agency of Research (AEI). Research at the CBMSO is facilitated by the Fundación Ramón Areces.

## Acknowledgements

We thank Julián Echave for interesting discussions, and Fernando Otero de Navascués for collaborating to a previous version of this work. We thank Centro de Supercomputación de Galicia (CESGA) for part of the computer resources.

## Notes

doi:10.5061/dryad.6wwpzgn2g

### Competing Interest Statement

The authors have declared no competing interest.

### Summary of Updates

We changed the definition of our structurally constrained models in such a way that the residue in the PDB at each site is not assigned deformation zero and, consequently, maximum fitness, but we compute the deformation by averaging over all possible initial residues. We corrected a mistake in the likelihood computation with raxml-ng, after which the log-likelihood of the empirical models with gamma distribution is higher than for our SSCPE models. We tested the SSCPE model for phylogenetic inference, comparing the maximum likelihood (ML) trees inferred with the SSCPE models and inferred with empirical models (with and without the options +F and +G) against trees inferred with structural divergences computed by the program PC_ali using the NJ and ME algorithms. The comparison of sequence-based and structure-based branch lengths detects accelerations of structural evolution possibly related to protein function changes. We introduced the new score "regularized maximum likelihood and minimal evolution" (REGMLAME) that unifies three widely used criteria for phylogenetic tree inference: maximum likelihood, minimal evolution and maximum posterior probability and reduces the biases inherent to the ML and ME mathods without normalization.

