## Supplementary Figures for "Site-specific structure and stability constrained substitution models improve phylogenetic inference"

### Supplementary Information for Site-specific structure and stability constrained substitution models improve phylogenetic inference

Ivan Lorca[1], Miguel Arenas[2,3] and Ugo Bastolla[1]
1 Centro de Biologia Molecular "Severo Ochoa", CSIC-UAM Cantoblanco, 28049 Madrid, Spain.
2 CINBIO (Biomedical Research Center), University of Vigo, 36310 Vigo, Spain.
3 Department of Biochemistry, Genetics and Immunology, University of Vigo, 36310 Vigo, Spain.


June 27, 2024

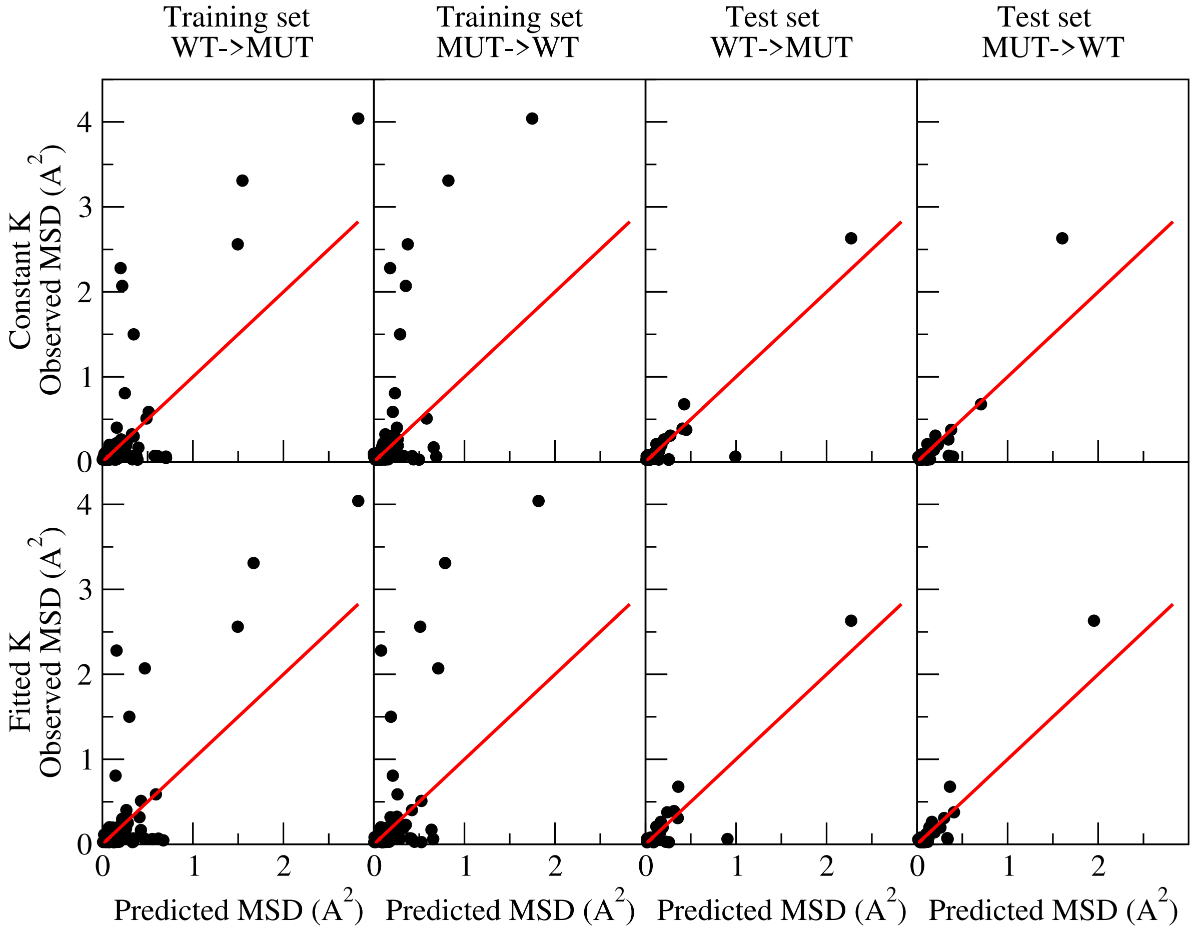


Figure S1: **Observed and predicted mean square deviation (MSD)** for the training set and the test set from wild-type to mutant and from mutant to wild-type, either keeping fixed the force constant of the torsional network model (upper line) or fitting it with B factors (lower line).

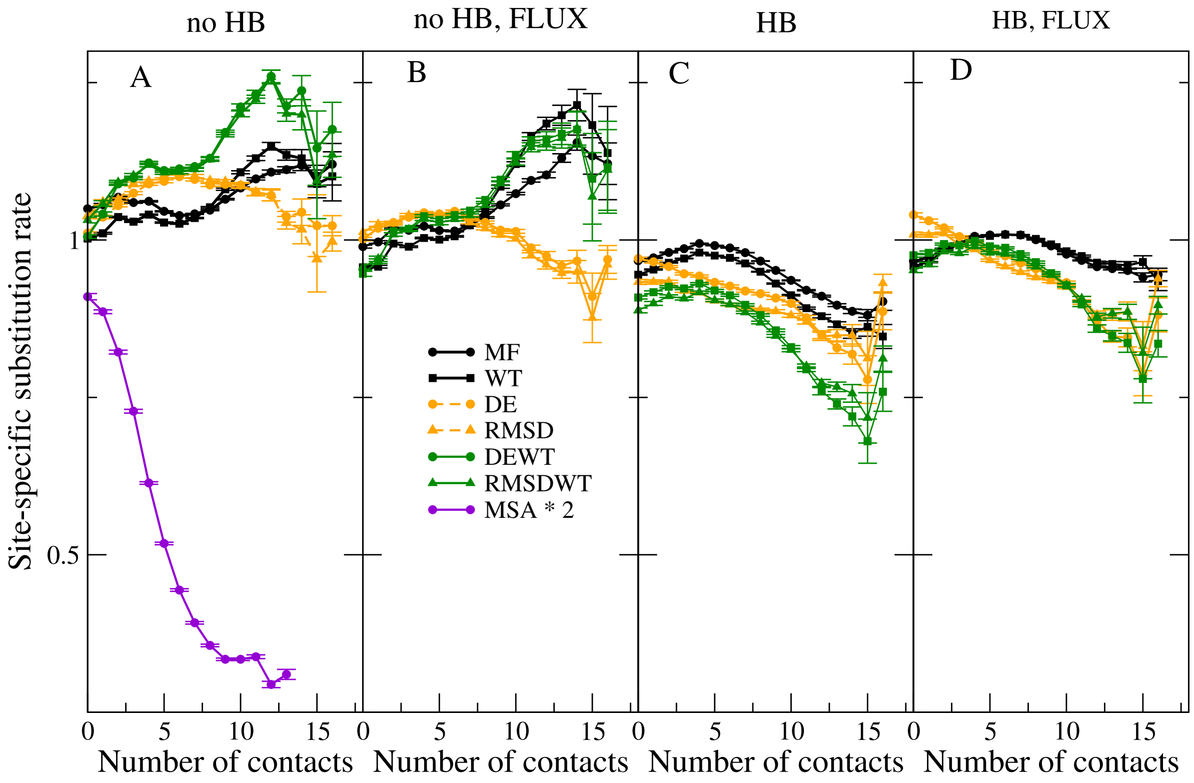


Figure S2: **Influence of different exchangeability models.** Average site-specific substitution rate as a function of the number of contacts of the site in the native state of the wild-type sequence (horizontal axis) for various SSCPE models. The exchangeability matrix at each site is computed either without (A,B) or with (C,D) the Halpern-Bruno model, either with the empirical exchangeability matrix (A,C) or with the matrix computed with the flux model (B,D). Data are averaged across all sites with the same number of contacts and 203 MSA.

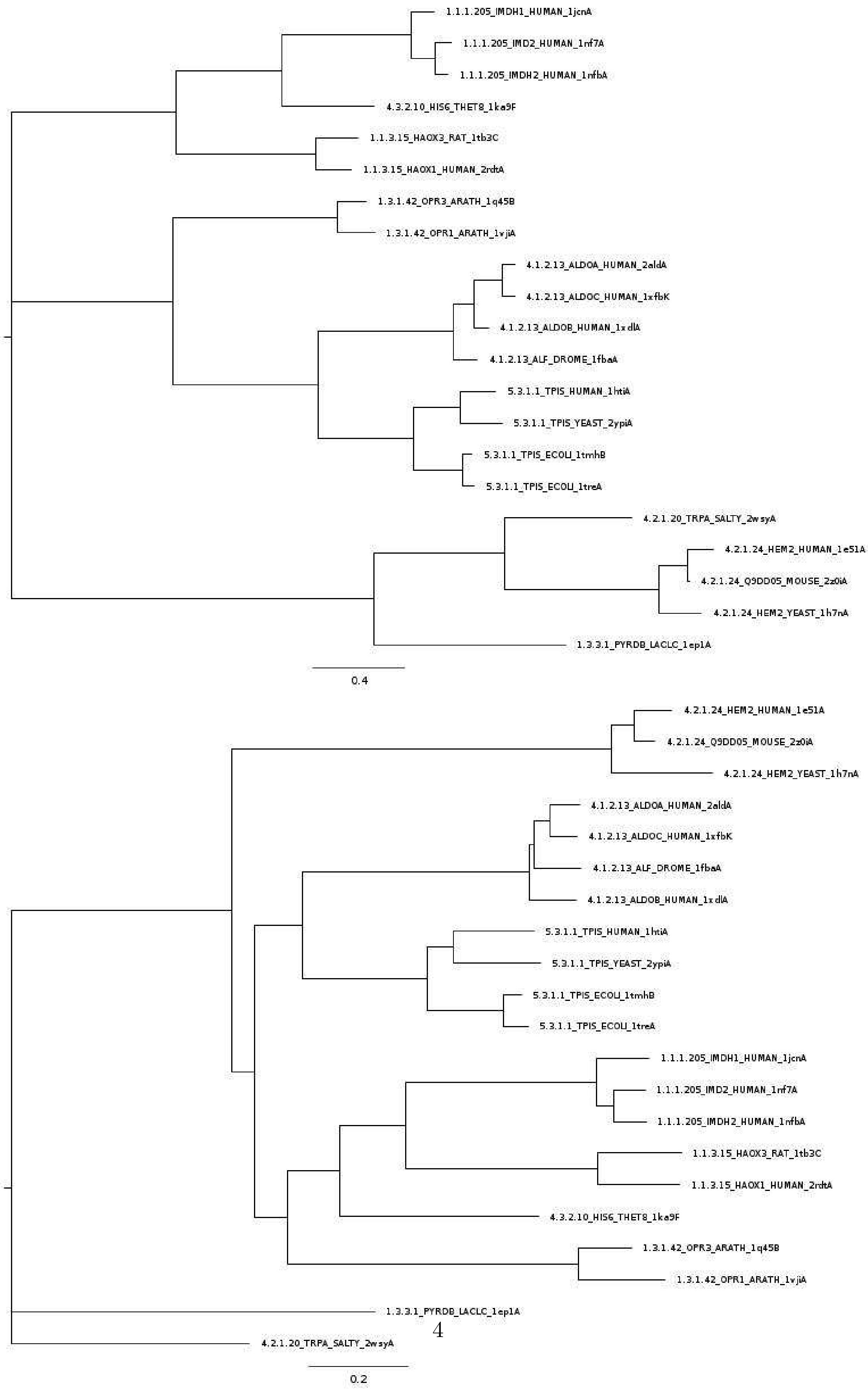




Figure S3: **Structure-based reference trees of the Aldolase superfamily.** Top: Neighbor Joining tree inferred with the program PC_ali. Bottom: Minimum Evolution tree inferred with the program FastME using the divergence matrix computed by the program PC_ali.

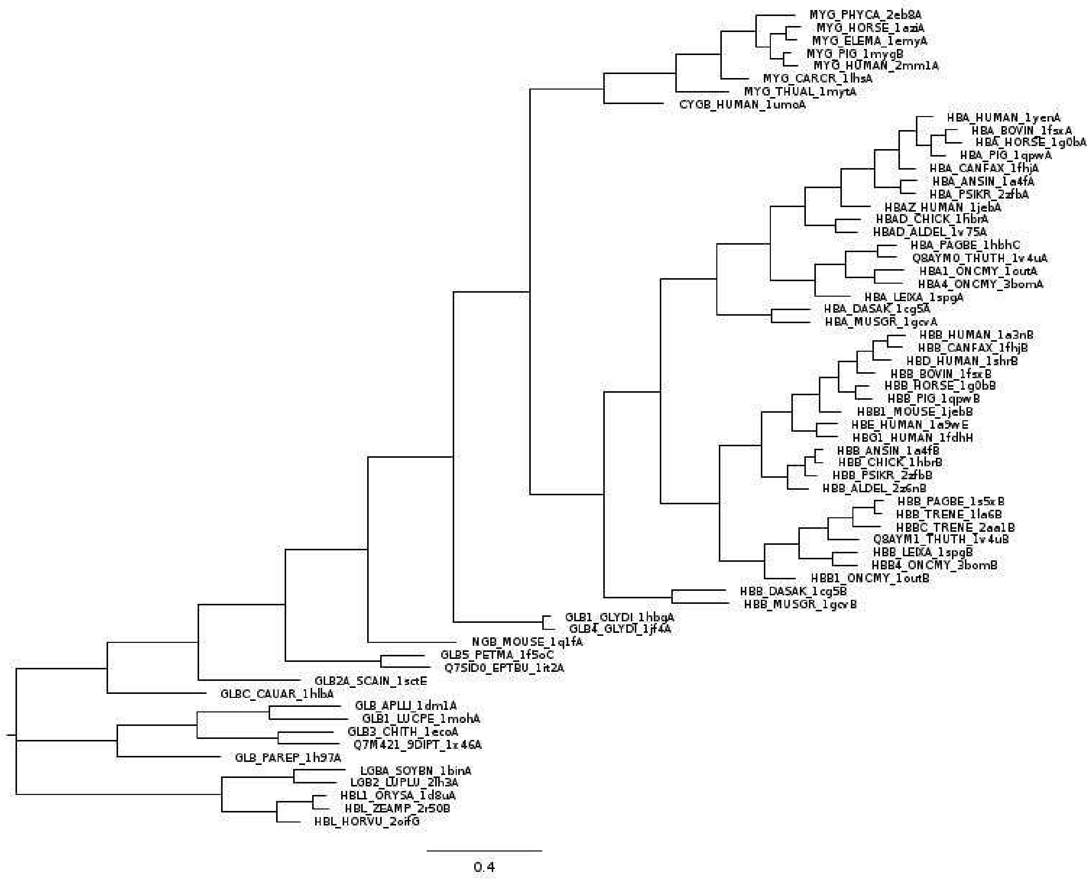


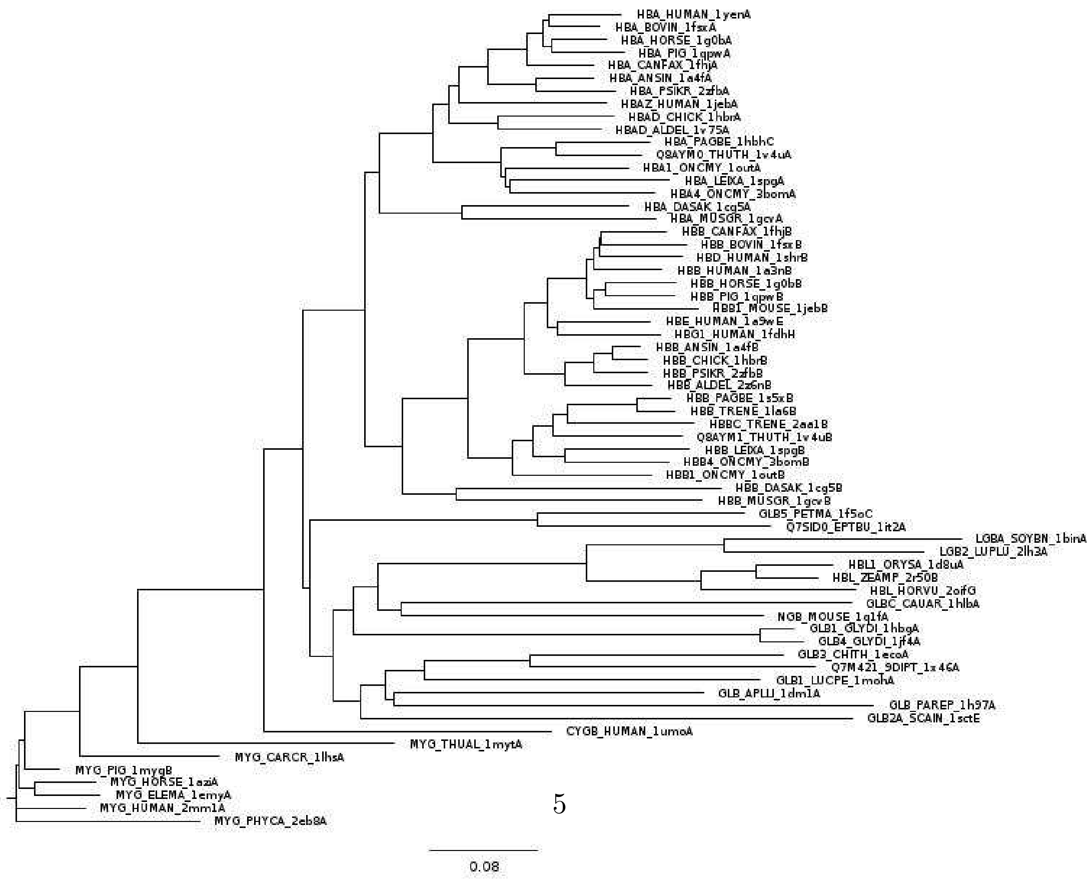




Figure S4: **Structure-based reference trees of the Globins superfamily.** Top: Neighbor Joining tree inferred with the program PC_ali. Bottom: Minimum Evolution tree inferred with the program FastME using the divergence matrix computed by the program PC_ali.

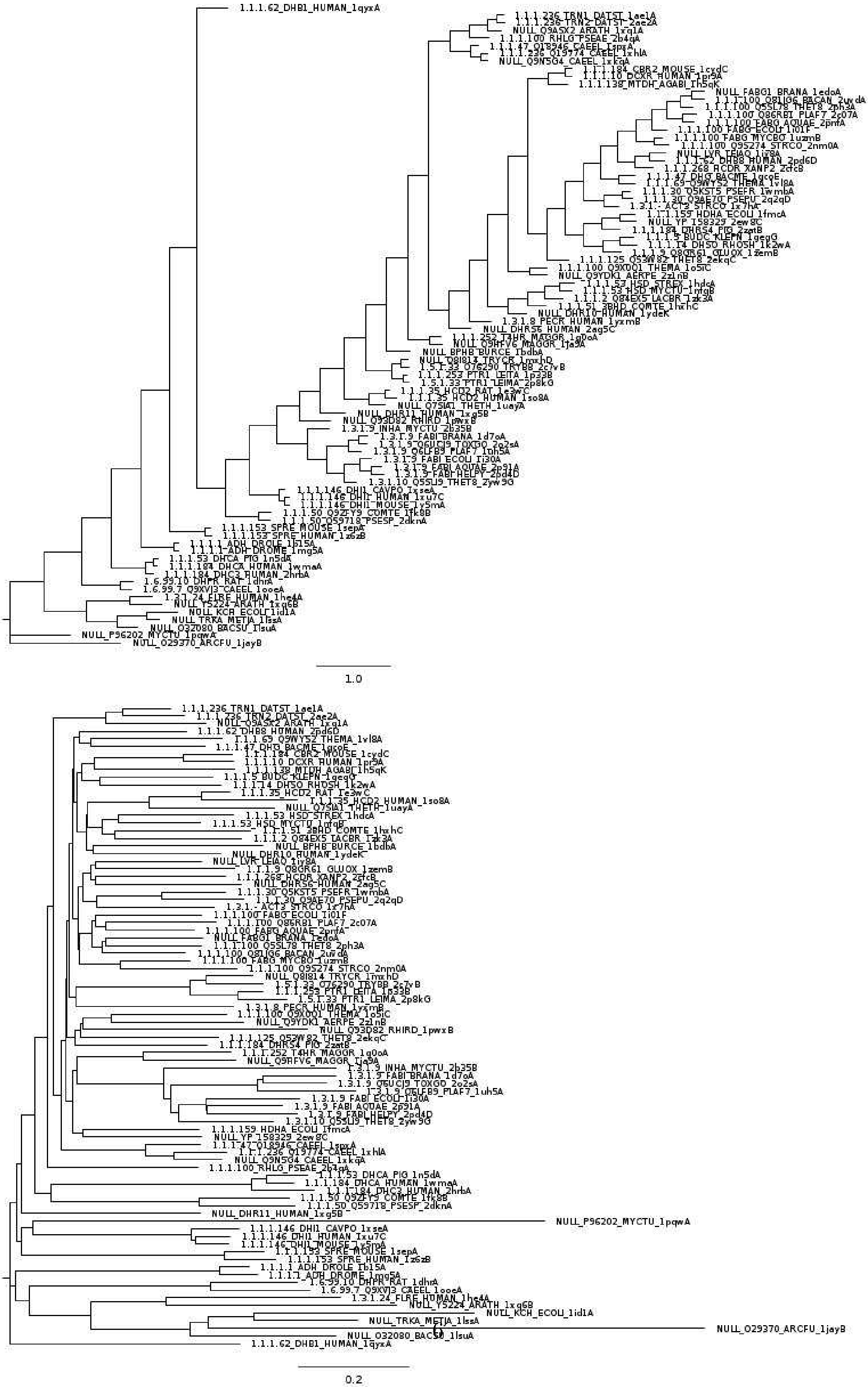


Figure S5: **Structure-based reference trees of the NADP superfamily.** Top: Neighbor Joining tree inferred with the program PC_ali. Bottom: Minimum Evolution tree inferred with the program FastME using the divergence matrix computed by the program PC_ali.

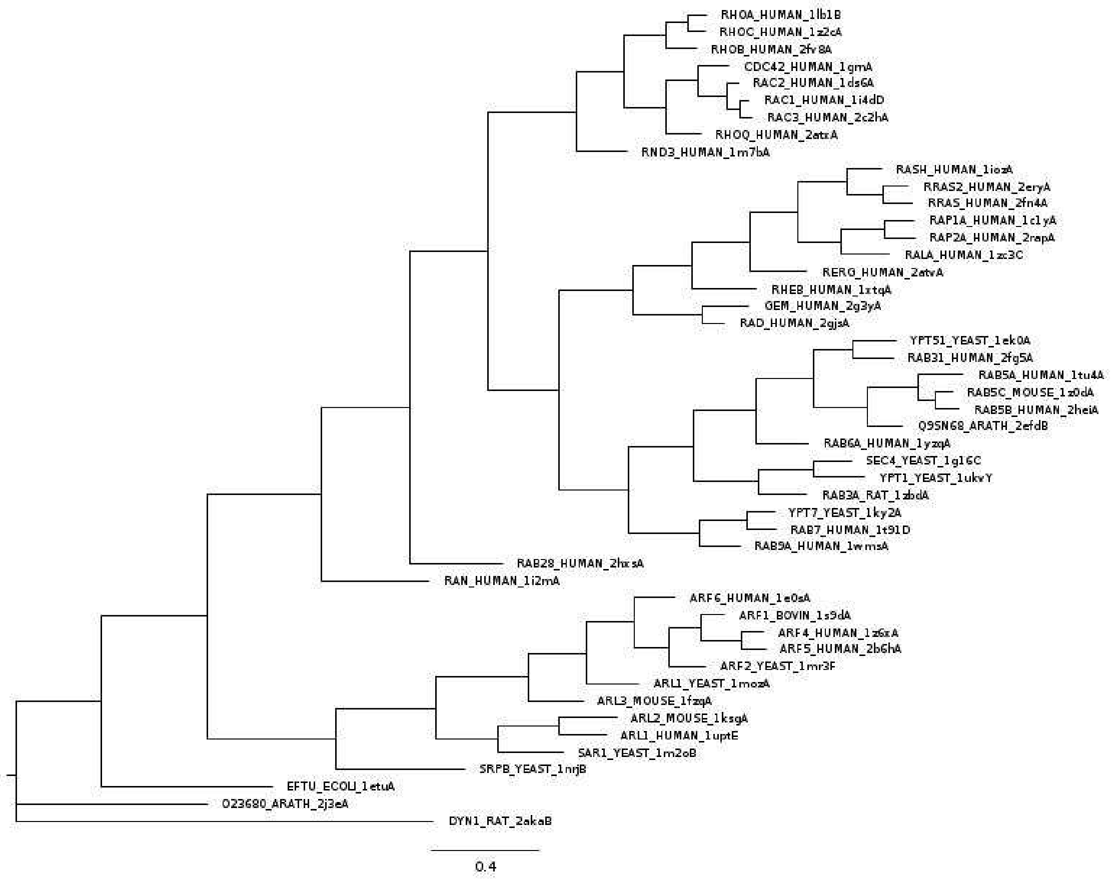


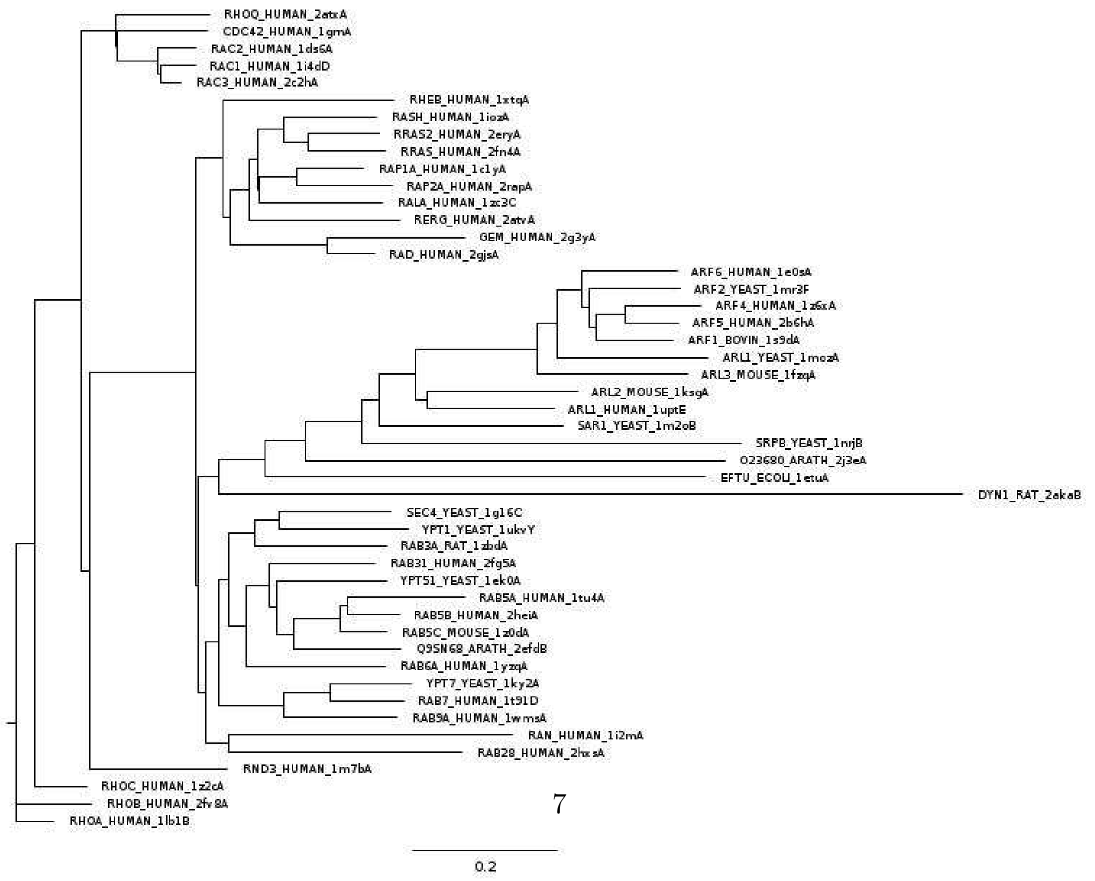




Figure S6: **Structure-based reference trees of the Ploop superfamily, cluster C1.** Top: Neighbor Joining tree inferred with the program PC_ali. Bottom: Minimum Evolution tree inferred with the program FastME using the divergence matrix computed by the program PC_ali.

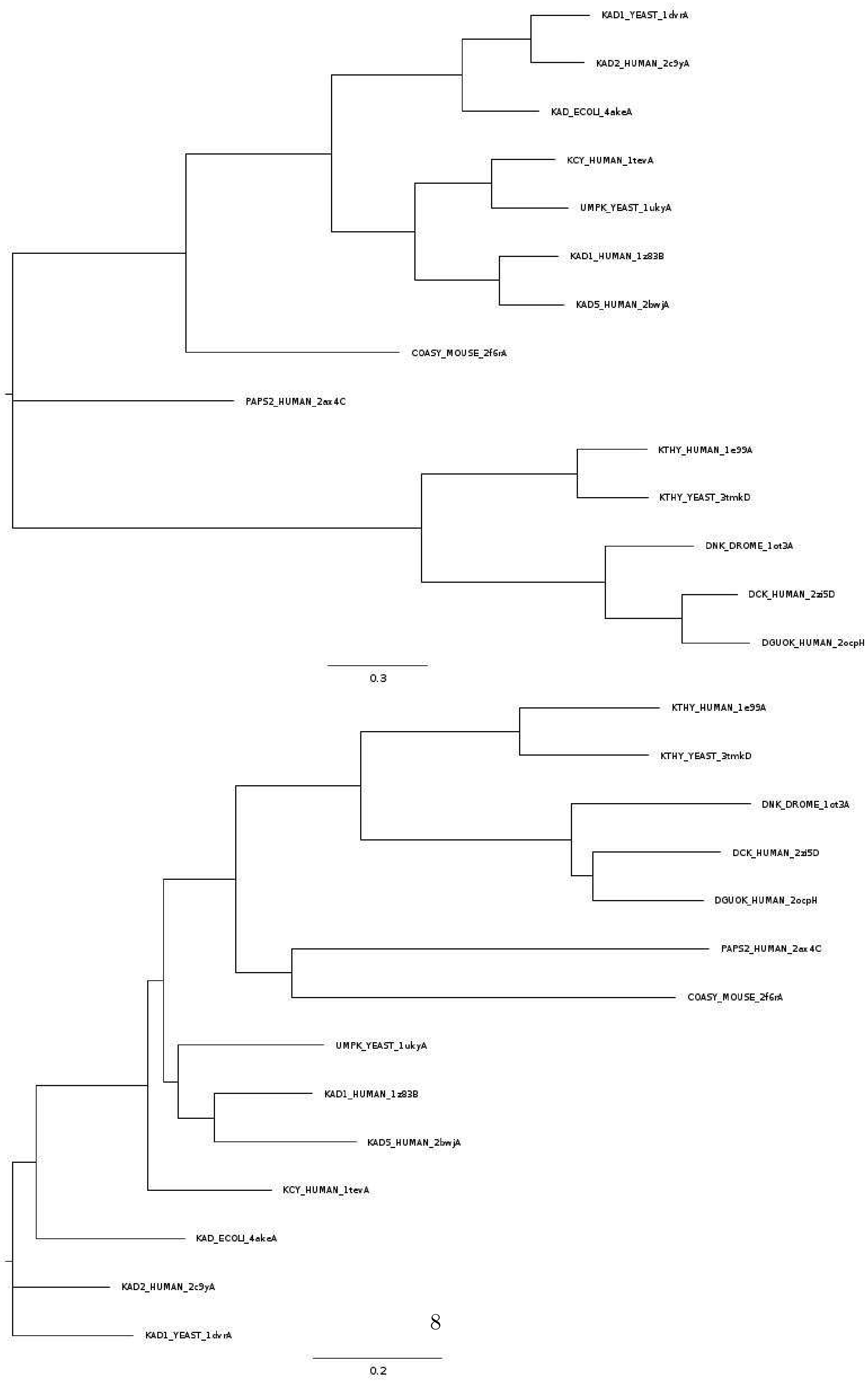




Figure S7: **Structure-based reference trees of the Ploop superfamily, cluster C2.** Top: Neighbor Joining tree inferred with the program PC_ali. Bottom: Minimum Evolution tree inferred with the program FastME using the divergence matrix computed by the program PC_ali.

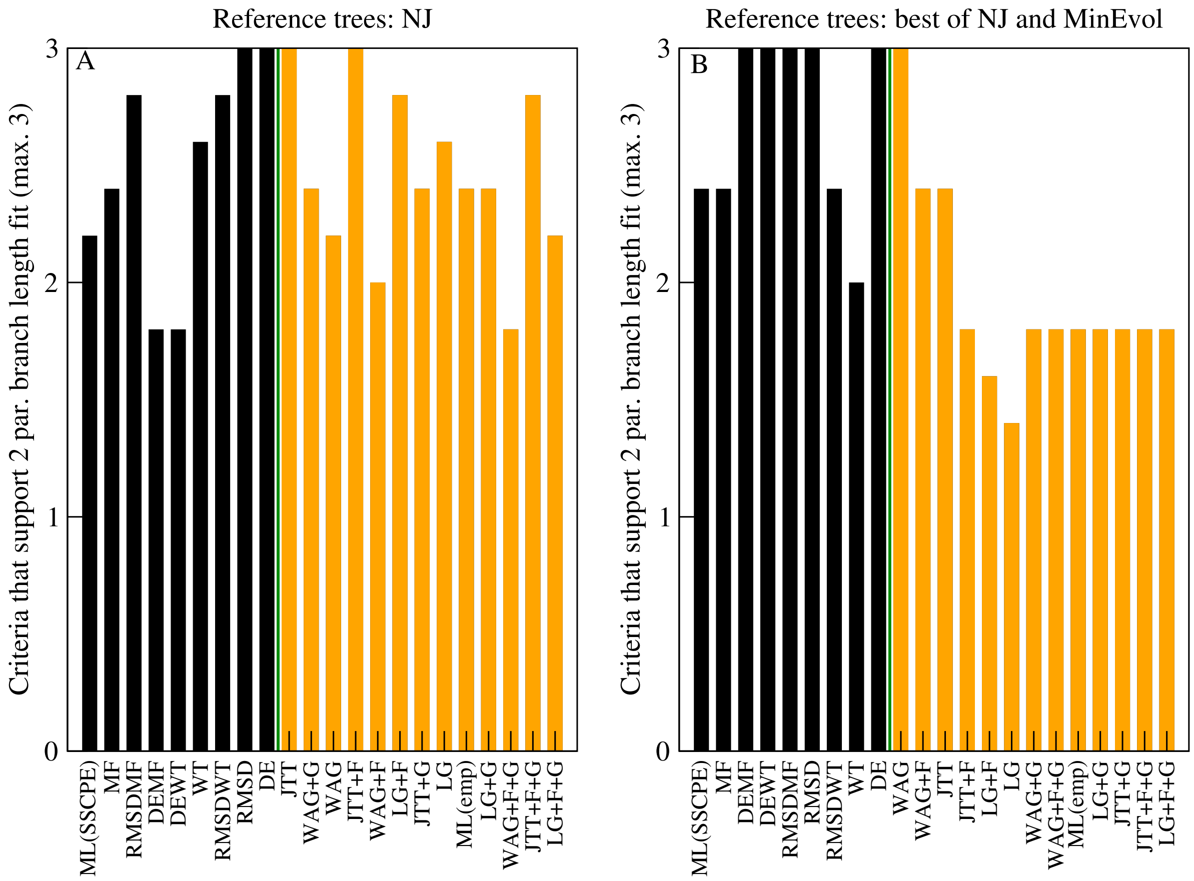


Figure S8: **Average number of criteria that support the 2-parameter fit of branch lengths against the 1-parameter fit.** The three tested criteria are AIC, corrected AIC and BIC. The substitution models are shown on the horizontal axis. Data are averaged over five superfamilies. A: Reference tree is NJ tree. B: Reference tree is selected with the REGMLAME criterion between the NJ and the ME tree.

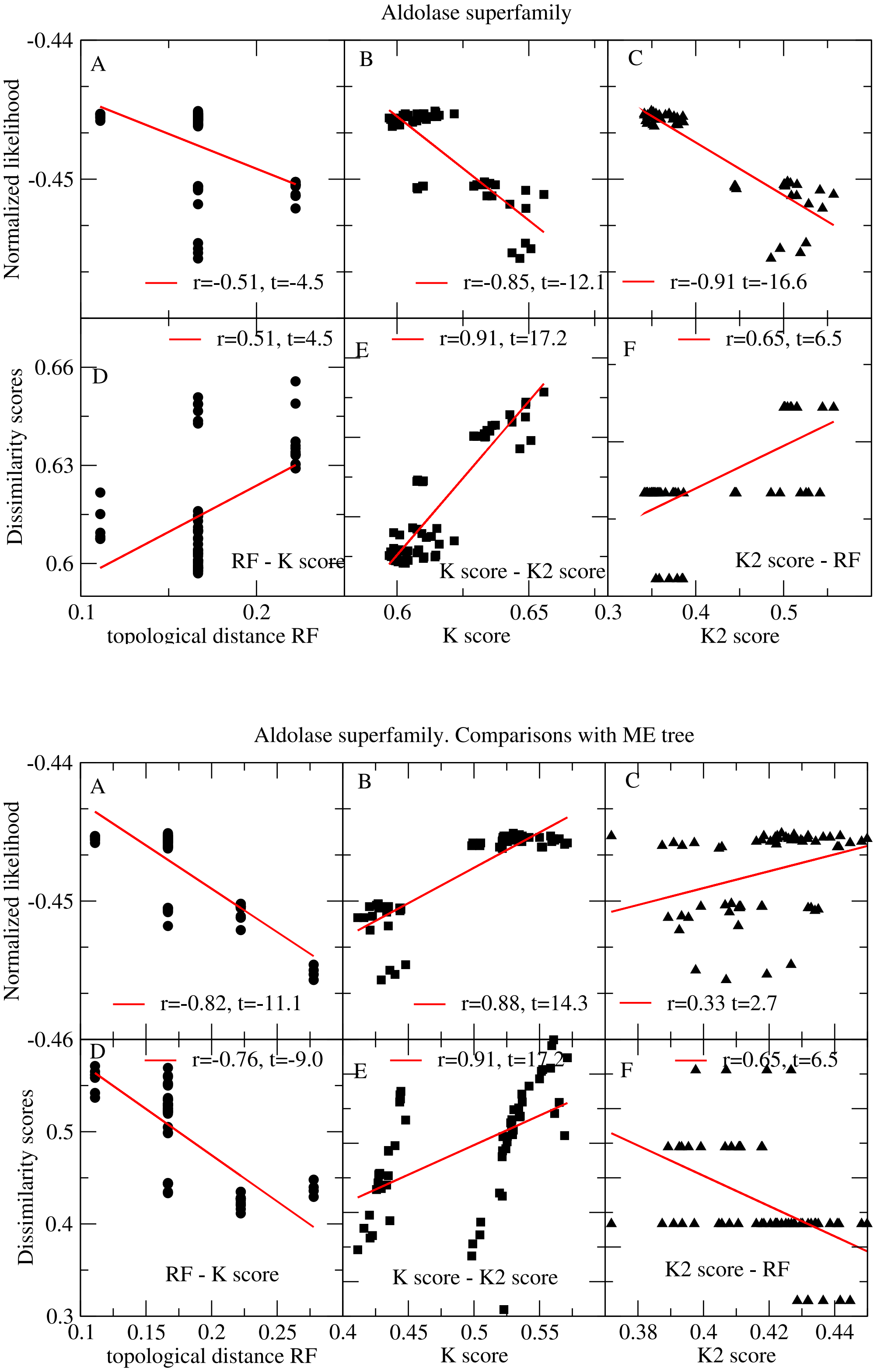


Figure S9: **Correlations between between dissimilarity from the reference tree of the Aldolase superfamily and log-likelihood scores.** Each point represents one of the ML trees of 12 empirical and 48 SSCPE substitution models. Above: NJ reference tree. Below: ME reference tree.

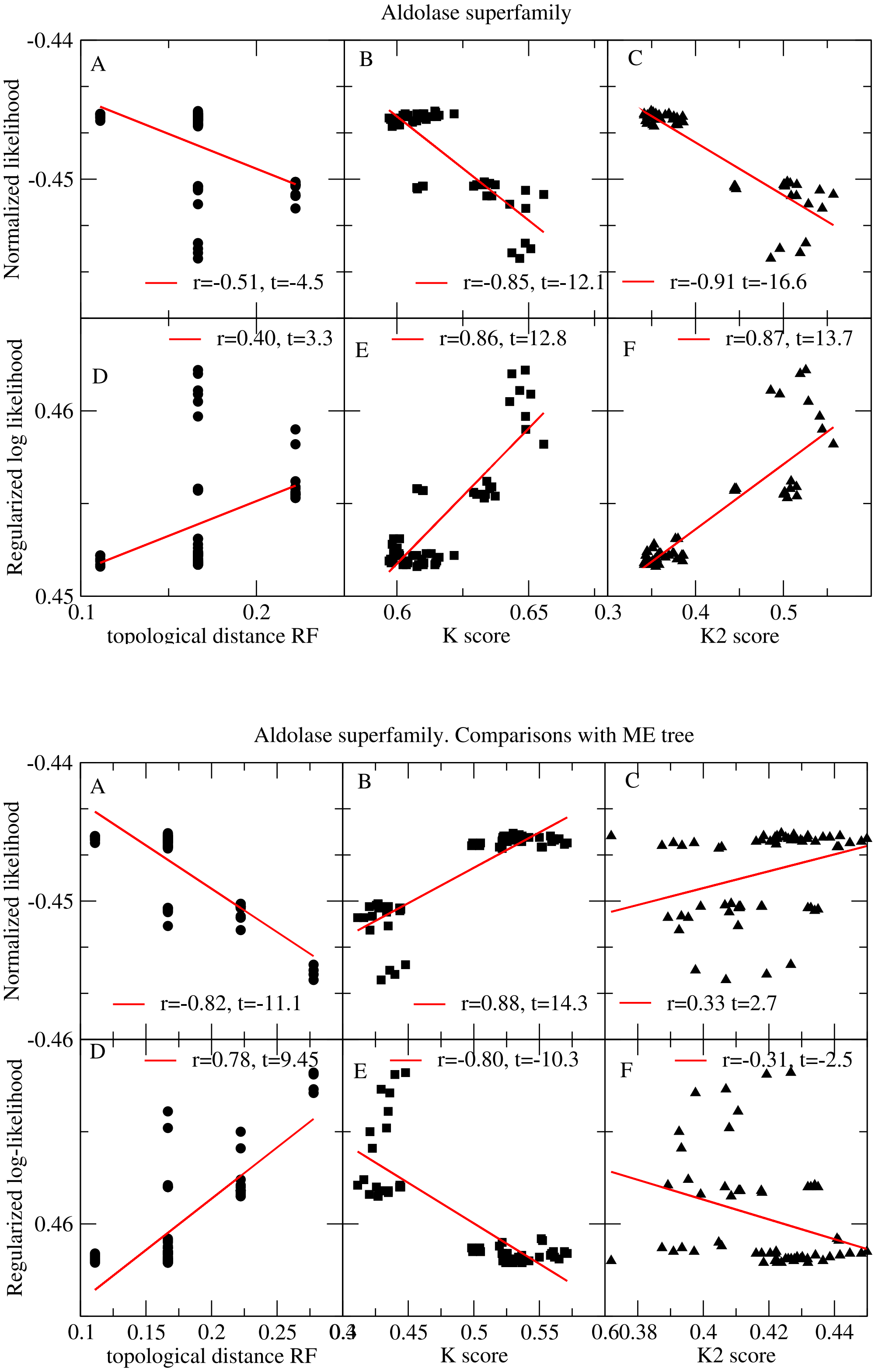


Figure S10: **Correlations between between differences from the reference tree of the Aldolase superfamily and regularized log-likelihood scores.** Each point represents one of the ML trees of 12 empirical and 48 SSCPE substitution models. Above: NJ reference tree. Below: ME reference tree.

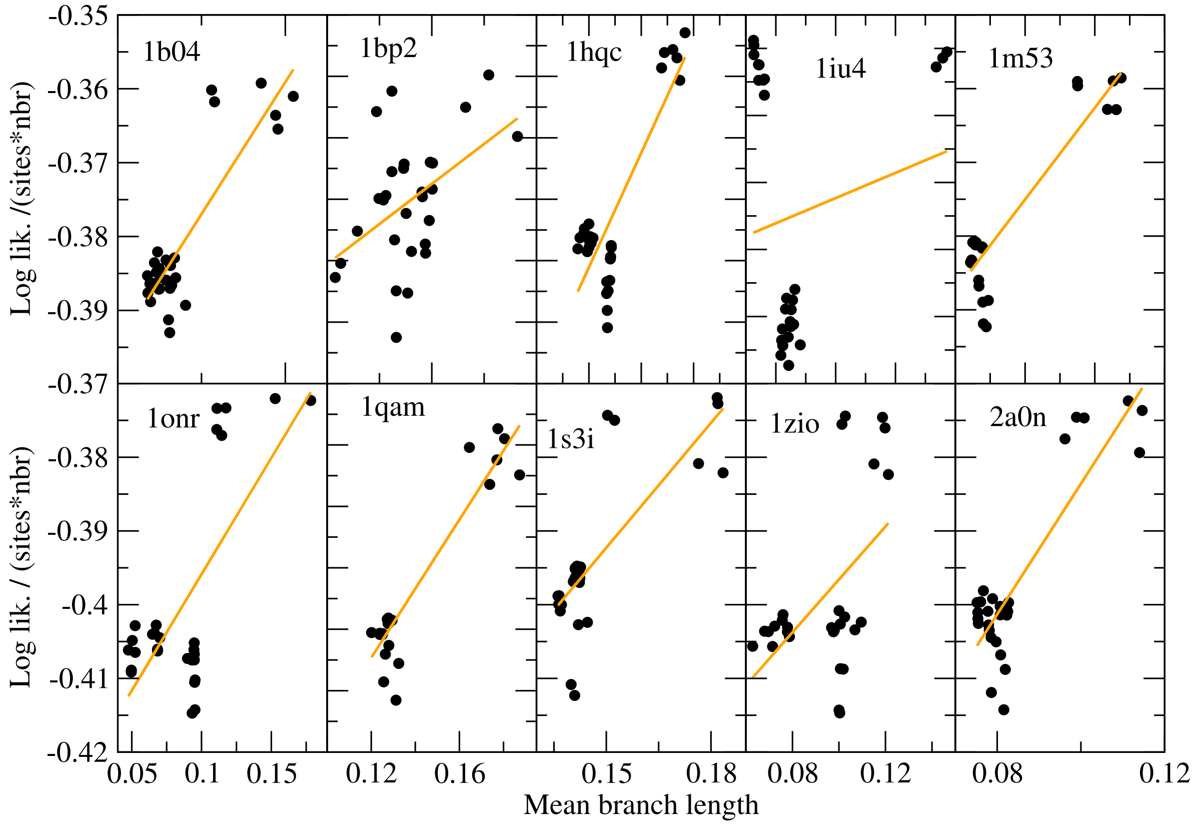


Figure S11: **Relationship between log-likelihood and branch length for ten large multiple sequence alignments of proteins with known experimental structure.** In each plot, points represent ML trees obtained with 28 different substitution models: 12 empirical substitution matrices (JTT, LG and WAG with all possible combinations of the options +F and +G) and the 8 SSCPE models with and without the option flux. The regularized fit of these scatter plots determined the parameters $\mu$ that we used for computing the regularized log-likelihoods of each model.

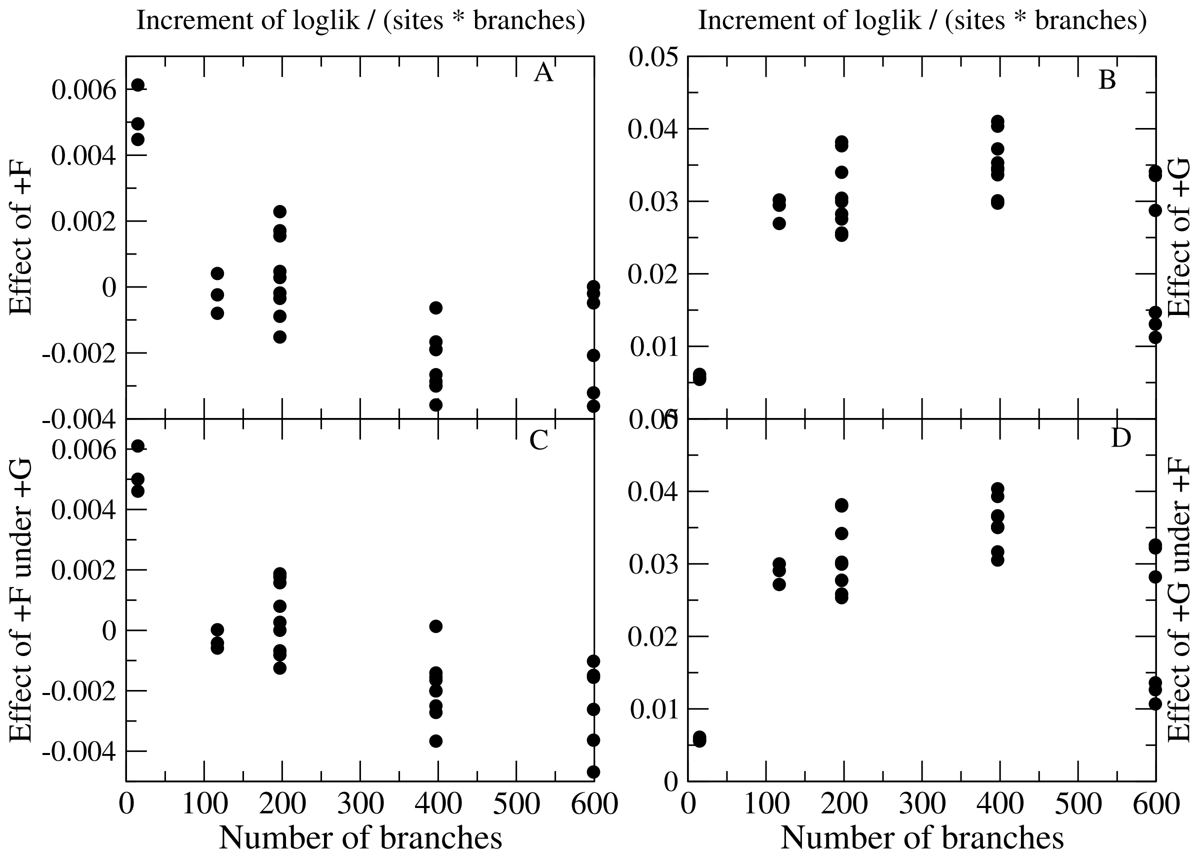


Figure S12: Increment of the log-likelihood per site and branch produced by the option +F with respect to the simple empirical model (A) and with respect to the empirical model with the option +G (C). Increment of the log-likelihood per site and branch produced by the option +G with respect to the simple empirical model (B) and with respect to the empirical model with the option +F (D).
